# Preconception Chronic Intermittent Ethanol Exposure Impacts Offspring Transcriptomes with Sex and Tissue Specific Effects

**DOI:** 10.64898/2026.06.29.735337

**Authors:** Rachel C. Rice, Richa R. Rathod, Daniela V. Gil, Remy R. Frawley, Laura B. Ferguson, Shirley Y. Hill, Gregg E. Homanics, Sean P. Farris

**Affiliations:** Center for Neuroscience at the University of Pittsburgh, Pittsburgh, PA, USA; Department of Anesthesiology & Perioperative Medicine, University of Pittsburgh, Pittsburgh, PA, USA; Bowles Center for Alcohol Studies, University of North Carolina at Chapel Hill, Chapel Hill, NC, USA; Department of Psychiatry, University of Pittsburgh, Pittsburgh, PA, USA; Department of Psychology, University of Pittsburgh, Pittsburgh, PA, USA; Department of Human Genetics, University of Pittsburgh, Pittsburgh, PA, USA; Department of Neurobiology, University of Pittsburgh, Pittsburgh, PA, USA; Department of Pharmacology & Chemical Biology, University of Pittsburgh, Pittsburgh, PA, USA; Department of Biomedical Informatics, University of Pittsburgh, Pittsburgh, PA, USA

## Abstract

Alcohol use disorder demonstrates ∼50% heritability, much of which remains unexplained by genetic sequence alone. Chronic alcohol exposure before conception changes offspring phenotypes through epigenetic mechanisms that are still being elucidated. Preconception ethanol exposure studies have focused on paternal exposure, neglecting maternal and biparental exposure. To address this, we exposed adult male and female mice to five cycles of chronic intermittent ethanol vapor interleaved with two bottle choice ethanol drinking and mated them to produce male and female F1 offspring with paternal, maternal, or biparental preconception ethanol exposure or controls. Whole blood and medial prefrontal cortex from adult, ethanol-naïve offspring underwent RNA-sequencing. We also analyzed previously unpublished RNA-sequencing data from male and female preimplantation embryos derived from preconception ethanol-exposed sires. Here, we report transcriptomic patterns of preconception ethanol exposure that depend on the exposed parent, offspring sex, and tissue which suggest metabolic and immune dysfunction in offspring.

## Introduction

Alcohol use disorder (AUD) is a complex disease hypothesized to arise from interactions between genetic predisposition and environmental influences.^1^ Genome-wide association studies have identified genetic risk variants that could contribute to AUD heritability,^2^ but a substantial amount of “missing heritability” of AUD remains unexplained by DNA sequence alone.^3^ Consideration of nongenomic and genomic inheritance may be essential to address missing heritability in AUD.^4^ While genomic inheritance relies on the transmission of DNA sequences to offspring, nongenomic inheritance involves environment-induced epigenetic changes affecting offspring gene expression.^4^ Thus, while genomic inheritance controls the *versions* of genes expressed in offspring, nongenomic inheritance determines *how* and *when* genes are expressed.

Preclinical studies of preconception ethanol exposure have begun to elucidate nongenomic inheritance mechanisms that could account for AUD missing heritability. Noncoding RNA (ncRNA) abundance in sperm has been implicated in the transmission of AUD- and anxiety-like behaviors from chronic ethanol-exposed sires to offspring.^5–7^ Paternal preconception ethanol exposure (PPE) also alters sperm histone H3 lysine 4 trimethylation in genomic regions regulating neurogenesis and craniofacial development.^8^ Rodent preconception ethanol exposure studies have also revealed sex-specific molecular and behavioral phenotypes in offspring, including developmental deficits and alterations in stress-, anxiety-like, and ethanol consumption behaviors.^4^

Investigating epigenetic heritability of preconception ethanol exposure phenotypes is critical given the prevalence of ethanol consumption by adults of reproductive age in the United States.^9, 10^ Preconception ethanol exposure remains understudied in comparison to prenatal exposure, although both produce deleterious effects persistent across generations.^4, 11–14^ Current preconception ethanol exposure literature additionally possesses significant sex bias and omission. Most published studies focus on PPE, with few investigating maternal or biparental preconception ethanol exposure (MPE and BPE, respectively), and few-to-none directly comparing sex-specific molecular outcomes in offspring.^4^ Studying PPE, MPE, and BPE in both offspring sexes enables a more comprehensive understanding of the molecular mechanisms and phenotypic outcomes of cross-generational ethanol exposure.

The current study examines the transcriptomic effects of chronic preconception ethanol exposure in the F1 generation, addressing the dearth of comprehensive research comparing PPE, MPE, and BPE outcomes in both sexes. We hypothesized PPE, MPE, and BPE produce sex-dependent transcriptional signatures in F1 blood and brain while altering shared pathways. To test this, we performed RNA-sequencing on whole blood and medial prefrontal cortex (mPFC) of adult, ethanol-naïve male and female F1 C57BL/6J mice following PPE, MPE, or BPE. We analyzed whole blood to identify potential biomarkers of preconception ethanol exposure. We selected mPFC based on prior rodent work showing germline-mediated changes in gene expression following chronic preconception stress or prenatal ethanol exposure (both fields which have informed preconception ethanol exposure research),^4, 15, 16^ as well as its dysfunction in decision making, impulse control, and associative learning in AUD pathogenesis.^17^ Thus, molecular mPFC alterations associated with preconception ethanol may indicate intergenerational AUD risk. We also analyzed previously-unpublished RNA-sequencing data from PPE preimplantation embryos to assess early developmental effects.^18^

In this work, we elucidate divergent transcriptomes of chronic preconception ethanol exposure dependent on parental exposure, offspring sex, and tissue analyzed. Despite this variation, we report gene expression changes in F1 with preconception ethanol exposure potentially indicative of metabolic and immune dysfunction. Furthermore, we find the rodent whole-blood transcriptome is a modest predictor of mPFC gene expression in this context. To our knowledge, the current study is the first comprehensive preclinical study of the transcriptomic outcomes of preconception ethanol exposure.

## Materials and Methods

### Animals

Adult (6 weeks of age at receipt) male (n=20 Air, n=21 CIEV) and female (n=18 Air, n=35 CIEV) C57BL/6J mice were acquired from The Jackson Laboratory (Bar Harbor, ME, USA; Strain #000664) and habituated to the University of Pittsburgh-specific pathogen free barrier facility for five days before experiments. Mice were maintained on a 12/12h light-dark cycle with *ad libitum* access to food and water and were provided a red plastic igloo for enrichment. Except for during 2BC (during which mice were single-housed), mice were group-housed with four (males) to five (females) cagemates. All experiments were approved by the Institutional Animal Care and Use Committee at the University of Pittsburgh and performed following the NIH Guide for the Care and Use of Laboratory Animals.

### Chronic Intermittent Ethanol Vapor-Two Bottle Choice

Founder generation (F0) mice underwent 5 cycles of CIEV-2BC lasting 11 weeks to simultaneously induce ethanol dependence and measure escalation of ethanol consumption.^19^ Mice were randomly assigned to ethanol exposure or control groups. Experiments were offset by one week between males and females (such that one sex was undergoing CIEV while the other was undergoing 2BC, **Supplementary Figure 1A**) to prevent potential confounding effects of sex-related pheromones on behavior from proximity in the vapor chambers, as filter tops were removed from cages to facilitate vapor inhalation. To ensure simultaneous completion of ethanol exposure before breeding, males underwent an additional round of 2BC while females completed a fifth CIEV cycle. Mice were group-housed during CIEV-exposure periods and single-housed for 2BC. To retain scent familiarity and prevent aggressive behavior in males, weekly group-housed bedding was retained from male cages and sprinkled into fresh cages during single-housing. Males were monitored after group housing for aggressive behavior warranting separation, which was not observed during experiments.

Cycles of continuous access 2BC lasted five overnight periods starting on Monday and ending on Saturday. On Monday, mice were weighed, single-housed, and provided with two 15-mL glass bottles with ball-bearing sipper tubes (Amuza, San Diego, CA, USA). Ethanol-exposed mice received one bottle with 15% (v/v) ethanol and one with water. Control mice received two bottles of water. Every 24h (∼4-5 h into the light cycle), bottles were weighed and their position was switched to mitigate side preference.

Fluid intake was measured in g/kg/day. Differences in ethanol consumption relative to baseline were assessed via a linear mixed effects model using the lmerTest (v. 3.1-3) and emmeans (v. 1.11.1) packages in R (v. 4.4.2).

CIEV exposure was performed as described previously,^18, 19^ lasting four overnight periods (starting on Monday) for 16h/day. On the first day of each CIEV cycle, mice were weighed. Each day of CIEV exposure, ethanol-exposed mice were given an intraperitoneal (*i.p.*) injection of 1.5 g/kg ethanol (Decon Labs, PA, USA) and 68 mg/kg of the alcohol dehydrogenase inhibitor pyrazole (Sigma-Aldrich, P56607-5G) in sterile saline. Control mice were given an *i.p.* injection of 68 mg/kg pyrazole in sterile saline. Mice were immediately placed into custom Plexiglass inhalation chambers following *i.p.* injection. Room air (8L/min) was introduced into two heated Erlenmeyer flasks, one of which received ethanol at a rate of 50-60 µL/min from a syringe pump (Harvard Apparatus, Holliston, MA, USA), before flowing into the vapor chambers. Ethanol content in the ethanol inhalation chamber was monitored by a custom sensor generously provided by Brian McCool (Wake Forest University). Following the final ethanol exposure each week, blood samples were collected from the tail vein using heparin-coated capillary tubes. Blood ethanol concentrations (BECs) were measured in plasma on an Analox Alcohol Analyzer (AM1, Analox Instruments, Lunenburg, MA, USA). Control mice also underwent blood collection for consistency of this stressor across groups, but this blood was not analyzed. Weekly inhalation chamber ethanol flow rate was adjusted based on the prior week’s BEC measurements to achieve pharmacologically relevant BECs of 100-200 mg/dL. After the final CIEV exposure during each cycle, mice underwent a 72h withdrawal period before 2BC.

### Breeding to Produce F1

Following a washout period from ethanol exposure with water (48h for males completing 2BC.5, 72h for females completing CIEV.5, **Supplementary Figure 1A**), F0 mice were randomly mated to produce PPE, MPE, BPE, or control offspring. Effects of PPE, MPE, and BPE on litter size and pup mortality relative to controls were assessed via linear mixed effects models in R. Group effects on litter sex ratios (measured as the proportion of female pups within litters) were assessed via a binomial generalized linear mixed model. Offspring born within a two-week period (n=8-9/sex/group) were selected to ensure a consistent age range. These F1 mice were weaned into groups in a randomized block design, such that each cage contained same-sex mice from each experimental group to control for the potential epigenetic group effects of behavior during rearing. F1 mice were raised to adulthood (6-8 weeks of age), kept ethanol-naïve, and remained undisturbed except for daily wellness checks and weekly cage changes.

### F1 Tissue Collection and RNA Sequencing

Whole blood was collected from adult, ethanol-naïve male and female F1 mice via terminal cardiac puncture. Mice were deeply anesthetized with isoflurane before a 25-gauge x 1 inch needle attached to a 3 mL tuberculin syringe was inserted into the heart. Blood (0.5-1 ml/mouse) was collected and 500 µL was immediately placed in an equal volume of DNA/RNA Shield™ (#R1200, Zymo Research, Orange, CA, USA). Mice were immediately decapitated and whole brain was collected and flash frozen in liquid nitrogen for storage at -80°C. Blood samples in DNA/RNA Shield™ were stored at 4°C for less than one week prior to RNA extraction.

Bilateral tissue punches from mPFC (defined as infralimbic, prelimbic, and cingulate cortex, area 1) were collected using a 1-mm biopsy punch (Rapid-Punch Sampling Kit, WellTech, Taiwan) from 100 µM sections taken in a Leica CM1950 Cryostat at -20°C, immediately placed on dry ice, and stored at -80°C until RNA extraction. Approximate anterior-posterior stereotaxic coordinates for mPFC tissue punches ranged Bregma +2.96 to +1.18. The Franklin and Paxinos Mouse Brain Atlas (Third Edition) was referenced to place tissue punches following neuroanatomical landmarks.

Whole-blood RNA was extracted using the Quick-RNA Whole Blood kit (#R1201, Zymo Research, Orange, CA, USA). Tissue punch RNA was extracted using the MagMAX™-96 Total RNA Isolation Kit. RNA concentration was quantified on a Qubit 4 Fluorometer using the Qubit RNA HS Assay Kit (ThermoFisher Scientific) and RNA integrity was analyzed using a TapeStation Instrument. Sample RNA concentrations averaged 97.1 ng/µL for whole blood and 24.4 ng/µL for mPFC. RNA integrity number equivalent values averaged 9.2 for whole blood and 9.7 for mPFC. Whole blood and mPFC RNA were submitted to the Genome Sequencing and Analysis Facility at the University of Texas at Austin for 3’Tag-Seq library preparation and RNA sequencing. Samples were sequenced on the NovaSeq S1 platform (Illumina, San Diego, CA) at a depth of 3-5 million paired end reads per sample (100 bp/read).

### PPE Embryo Study

Male and female PPE embryo RNA-sequencing data were collected during a prior PPE study in our laboratory^18^ but were not included in the prior publication of this work. In this study, 9-10-week-old male C57BL/6J mice purchased from The Jackson Laboratory (Bar Harbor, ME, USA) were randomly assigned to CIEV or control groups and underwent CIEV for 16h/day, 4 overnight periods/week (starting Monday) for six weeks and for three overnight periods (starting Monday) the 7^th^ week. The CIEV exposure protocol followed the methods described above for blood and brain study, except 2BC and single-housing were excluded. Immediately after the final CIEV exposure on the 7^th^ week, control and PPE males were mated to ethanol-naïve, super-ovulated female C57BL/6J mice. The following morning, single-cell embryos were collected from oviducts and cultured in KSOM mouse embryo media (Millipore Sigma #MR-106-D, Darmstadt, Germany) until the blastocyst stage (3.5 days post conception).

Blastocysts were prepared for library preparation and sequencing by placing in acidic Tyrode’s Solution (Millipore Sigma #T1788) to remove the zona pellucida, washing with M2 medium (Millipore Sigma #M7167), and placing in lysis buffer (2.5mM dNTPs, 1 U/µl RNase Inhibitor, 0.1% Triton X-100) for storage at -80°C. RNA concentrations of samples used for RNA-sequencing ranged 12.4-23.6 ng/µL. A total of 24 samples (n=5-7/sex/group) passed quality control, which includes RNA integrity values > 7. Samples were sequenced by BGI (Shenzhen, China) on the BGISEQ-500 platform at a depth of 80-130 million paired-end reads per sample (100 bp/read). A total of n=xx/sex/group embryo samples were bioinformatically analyzed.

### Bioinformatic Analyses

For all samples (blood, mPFC, and embryo data), FASTQC (v. .0.11.9) was used to assess read quality. A total of n= 8-9/sex/group blood, n=4-8/sex/group mPFC, and n=5-7/sex/group embryo samples passed quality control and were bioinformatically analyzed. Adapters were trimmed from paired end reads using CutAdapt (v. 3.5), and bwa (v. 0.7.17) was used to map reads to the mouse genome (GRCmm39 assembly). Raw counts were quantified from mapped reads using HTSeq (v. 2.0.5). Trimmed reads and raw count matrices from this study have been deposited to the Gene Expression Omnibus under accession number GSE329570. Following raw count matrix generation, all analyses were performed in R. Embryonic sex was determined through principal component analysis on counts per million (CPM) expression values of the Y chromosome genes *Ddx3y*, *Kdm5d*, and *Eif2s3y* (**Supplementary Figure S2**). Embryonic samples formed two distinct clusters indicating genetic sex; Samples with appreciable expression of *Ddx3y*, *Kdm5d,* and *Eif2s3y* were identified as male, whereas samples with undetected (<1 CPM) *Ddx3y*, *Kdm5d*, and *Eif2s3y* were identified as female.

Differential expression analysis was performed using the Bioconductor package edgeR (v. 4.6.2), comparing PPE, MPE, and BPE to control samples within sexes and tissues. Differentially expressed genes (DEGs) were defined as those with p_nominal_ < 0.05 and |log_2_FC| > 1.5 to balance type I and type II error rates while allowing for sufficient DEG numbers for functional enrichment analyses. Within each group/sex/tissue combination, DEGs were assessed for functional enrichment using the web-based applications Enrichr (accessed through its R interface, enrichR) referencing the Reactome 2024 database and Ingenuity Pathway Analysis (IPA; QIAGEN, Inc., Aarhus, Denmark).

To comprehensively assess shared pathways affected by intergenerational preconception ethanol exposure, we performed a robust gene set enrichment analysis (GSEA) on DEGs using the clusterProfiler (v. 4.16.0) R package. We referenced all 238 gene ontology and functional enrichment databases available through Enrichr. For each group/sex/tissue combination, the GSEA function was run using the fgsea method, with a minimum gene set size of 15 and a maximum gene set size of 500. Default settings were utilized for all other parameters. Overlaps in DEGs as well as significantly (p_nominal_ < 0.05) enriched pathways across sexes, groups, and tissues were visualized using the VennDiagram (v. 1.7.3) and UpSetR (v. 1.4.0) packages.

Although thresholding results by p-value and |log_2_FC| cutoffs can illuminate the most dramatic changes in gene expression, this results in a trade-off between statistical and biological relevance.^20^ Thus, we utilized the improved rank-rank hypergeometric overlap (RRHO2) algorithm developed by Cahill *et al.*^20^ to assess concordant and discordant gene expression patterns between the sexes, within groups and tissues. This algorithm ranks genes from two datasets based on their associated p-value and fold change expression values and can detect statistically significant, biologically relevant overlaps in gene expression that are missed with thresholding.^20^

Finally, to determine whether the whole-blood transcriptome could predict mPFC gene expression and/or preconception ethanol exposure status, we followed the methods described in Ferguson *et al.* (2022)^21^ to: (1) Perform between- and within-subjects gene expression correlations across blood and brain and (2) Utilize machine learning to classify samples as preconception ethanol-exposed or controls based on whole blood gene expression. We separated both analyses by sex for consistency with Ferguson *et al.* (2022)^21^ and due to the unique gene expression signatures between males and females observed in our differential expression and RRHO2 analyses. PPE, MPE, and BPE were collapsed into a single preconception ethanol group to prevent overfitting. For both between- and within-subjects correlations, we filtered out lowly-expressed genes (genes were retained if CPM exceeded 1 in at least 8 samples), applied TMM normalization, and extracted voom-normalized expression values for each gene in blood and mPFC for each subject. For the between-subjects correlations, we averaged the normalized expression of each gene across subjects within sex and calculated the Spearman ρ correlation coefficient for mPFC versus blood gene expression using the corr.test function from the psych (v. 2.5.6) R package. For within-subjects analyses, we extracted the normalized gene expression in mPFC versus blood for each subject, then computed Spearman ρ across subjects, and correlated blood versus mPFC expression using paired values from each subject. P-values were corrected for multiple comparisons using the Holm–Bonferroni method, and correlations with p_adj_ < 0.05 were considered significant. To determine genes that correlate between blood and mPFC across generations (as Ferguson et *al.* examined the F0 generation following CIEV^21^), we compared genes with significant within-subjects correlations from our study to those from the blood-mPFC within-subjects correlations performed by Ferguson *et al.* (2022).^21^

We used four different machine learning classification models to discriminate subjects’ preconception-ethanol status based on blood gene expression. The first three models were identical to those used in Ferguson *et al.* (2022).^21^ These included logistic regression with elastic net regularization (LR), random forest (RF), and partial least squares discriminant analysis (PLSDA) models within the MLSeq (v. 2.28.0) R package. In addition, we used the extreme gradient boosting (XGB) model within the caret (v. 7.0-1) R package due to its flexibility of use with high-dimensional data with lower sample sizes. Like Ferguson *et al.* (2022),^21^ we utilized five-fold cross-validation repeated ten times to choose the optimal hyperparameters for each model. For male samples, the optimal value for the RF hyperparameter mtry was determined to be 2. The elastic net model selected α = 0.2125 and λ = 0.2118. The optimal number of components for PLSDA was ncomp = 3. For XGBoost, the optimal hyperparameters were: nrounds = 100, max_depth = 2, eta = 0.4, gamma = 0, colsample_bytree = 0.8, min_child_weight = 1, and subsample = 0.556. For females, the optimal RF mtry hyperparameter value was determined to be 10,445, the elastic net model selected α = 1 and λ = 0.251, and the PLSDA model selected ncomp = 3. For XGBoost, the optimal hyperparameters were: nrounds = 50, max_depth = 4, eta = 0.4, gamma = 0, colsample_bytree = 0.8, min_child_weight = 1, and subsample = 1. As in Ferguson *et al.* (2022),^21^ we evaluated the performance of the classifiers by calculating the area under the receiver operating characteristic curve (AUC) using the roc function from the pROC package (v. 1.19.0.1). We also extracted the most important features (*i.e.,* genes) for discriminating between samples with preconception ethanol exposure and controls using the varImp function from the caret package.

## Results

### F0 Chronic Intermittent Ethanol Vapor Exposure & Breeding

To generate PPE, MPE, BPE, and control F1 offspring, we exposed male and female founder (F0) mice to five cycles of interleaved chronic intermittent ethanol vapor exposure (CIEV) and two-bottle choice (2BC; **Methods, Supplementary Figure S1A**). CIEV exposure drove increased voluntary ethanol consumption in both sexes relative to baseline (p < 0.001), an effect observed after the first CIEV cycle (**Supplementary Figure 1B**). Consistent with prior studies, females drank more ethanol than males during all cycles (p < 0.001; **Supplementary Figure S1B**). Pharmacologically-relevant blood ethanol concentrations (BECs) averaging 150-200 mg/dL^19^ were recorded throughout CIEV and did not differ by sex (**Supplementary Figure S1C**; p = 0.14). There were no differences in litter sex ratios between groups (**Supplementary Table S1**). BPE mice produced ∼1.3 fewer pups relative to controls (p = 0.04), though this difference was not significant when adjusted for multiple comparisons (Tukey p_adj_ = 0.15, **Supplementary Table S1**). MPE increased pup mortality relative to controls, with ∼1.4 more pups per litter dying before postnatal day three (p = 0.01; Tukey p_adj_ = 0.042; **Supplementary Table S1**).

### Whole-Blood Differential Gene Expression

We raised male and female PPE, MPE, and BPE F1 mice to adulthood, keeping them ethanol-naïve with minimal disturbance before sacrifice at 6-8 weeks of age (**Methods**). 3’Tag-Seq on RNA extracted from blood detected 22,756 genes with nonzero counts across groups (**Supplementary Table S2**), along with varying patterns of differentially expressed genes (DEGs) by group and sex at a threshold of |log_2_FC| > 1.5 and p_nominal_ < 0.05. Across groups and sexes, protein-coding RNAs represented the most DEGs (∼75%-90%), followed by long noncoding RNAs (lncRNAs; ∼7-13%), pseudogenes (∼2-10%), and various small ncRNAs or RNAs with biotypes yet to be experimentally confirmed (TEC; <1%; **Table 1**). PPE male offspring had more upregulated DEGs, whereas PPE female DEGs were preferentially downregulated (**Table 1**; **Figure 1A**). MPE elicited the largest shift in blood transcriptomic profiles, producing the most DEGs in both sexes with predominantly upregulated expression (**Table 1**; **Figure 1A**). BPE male and female DEGs were near-evenly split between up- and downregulation (**Table 1**; **Figure 1A**).

**Figure 1.**
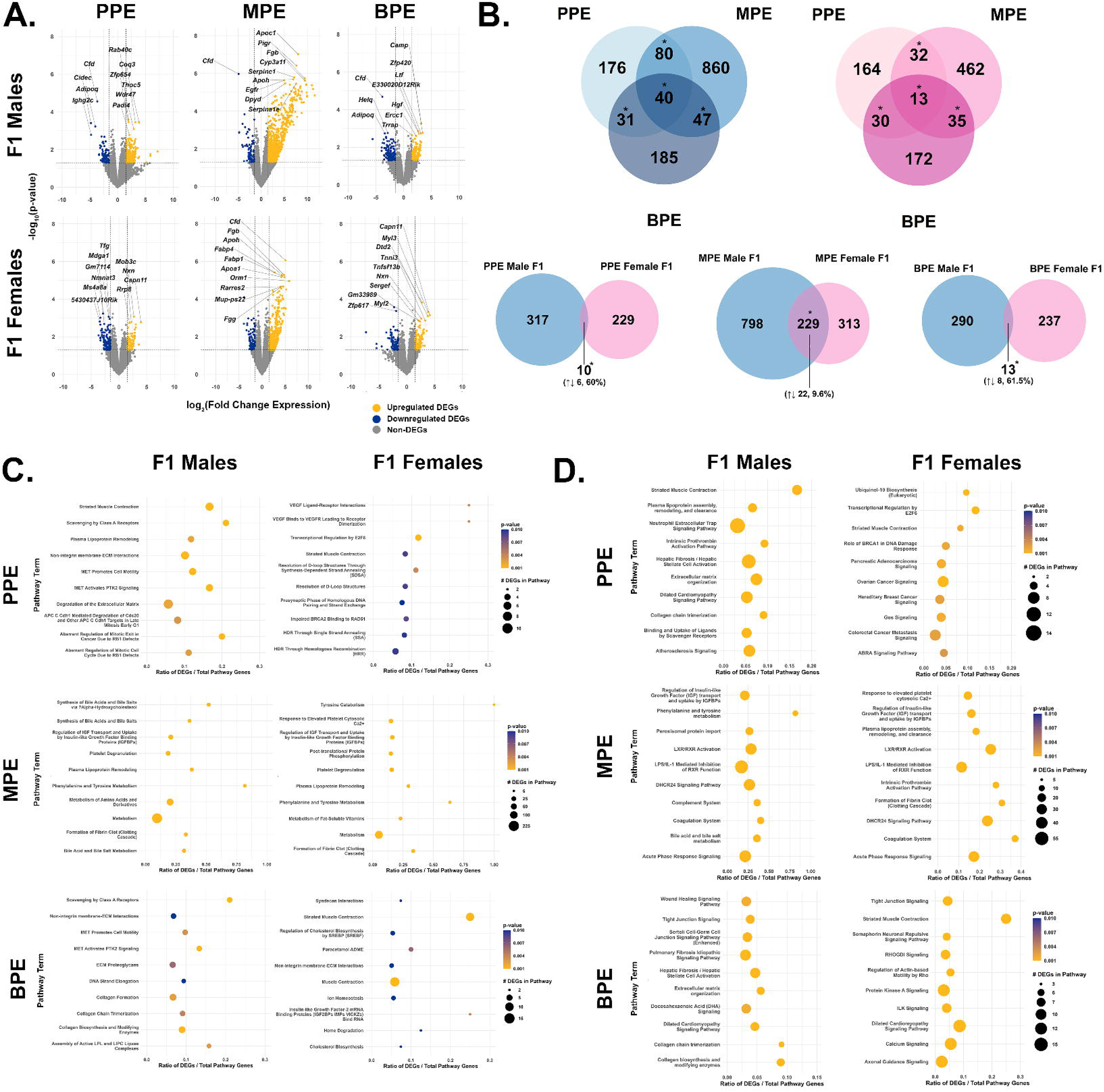
Adult F1 whole blood displays unique differential gene expression signatures based on sex and mode of preconception ethanol exposure. (A.) Volcano plots highlighting differentially expressed genes (DEGs), organized by sex and exposure group. DEG thresholds of p < 0.05 and |log_2_FC| > 1.5 are indicated with dotted lines. (B) Venn diagrams comparing DEGs: (top) across exposure groups, within sexes and (bottom) between males and females within exposure groups. All overlaps in gene expression indicated with * were significant by χ² analysis, p < 0.01. (C.) DEG Functional enrichment analysis results from Enrichr utilizing the Reactome 2024 database, organized by sex and exposure group. Top ten terms by p-value are displayed. (D.) DEG functional enrichment analysis results from Ingenuity Pathway Analysis, organized by sex and exposure group. In (C.) and (D.), p-values were truncated at p = 0.001 for ease of display and to ensure a consistent color gradient between males and females.

**Table 1.**
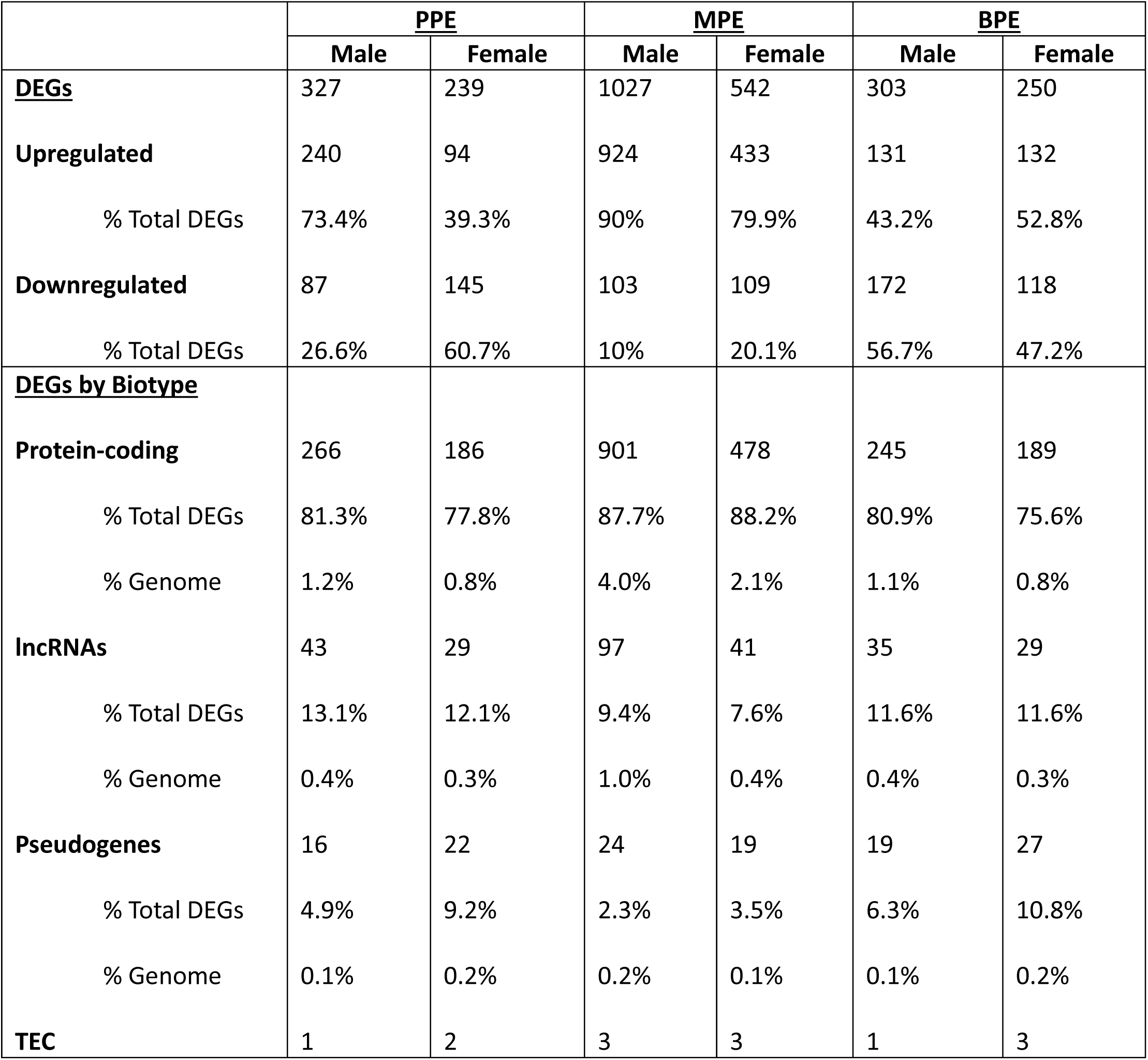
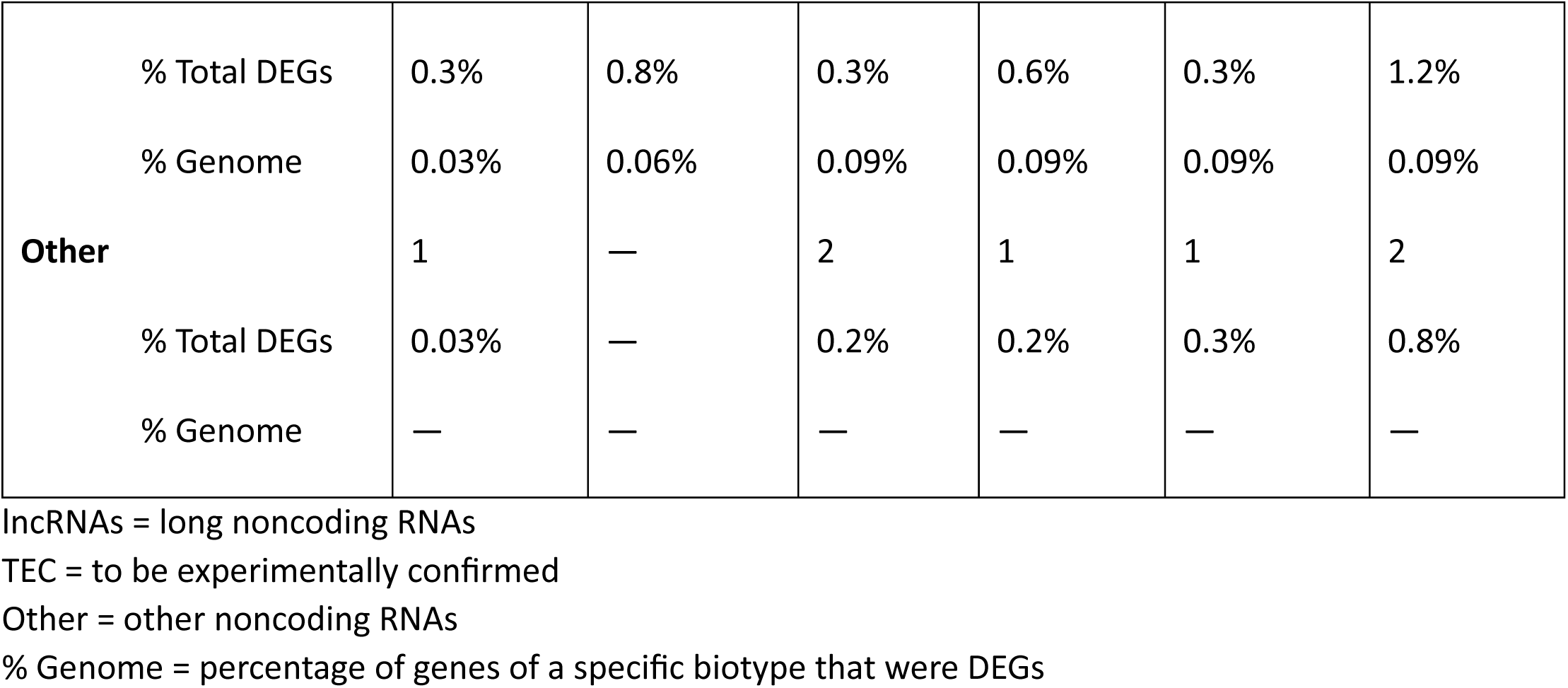
Whole blood differential gene expression.

Top DEGs by p_nominal_ across groups included genes encoding immune-related peptides, complement factors and their receptors, clotting factors, lipo- and glycoproteins, and several enzymes (**Figure 1A**).

Although Pearson’s Chi-squared tests with Yates’ correction were significant between groups/within sexes, shared DEGs were few. Forty DEGs were shared across groups in males (**Figure 1B**; **Supplementary Table S2**), χ²(3, N = 21,594 detected genes in males) = 1866.6 , p < 0.001. These included *complement factor D* (*Cfd*)*, cell death-inducing DFFA-like effector C* (*Cidec*)*, adiponectin* (*Adipoq*), and *fatty acid binding protein 4* (*Fabp4*), which were downregulated across groups (**Supplementary Table S2**). Females similarly had small, significant overlaps in DEGs between groups (**Figure 1B**). Only 13 DEGs were shared across all female groups, χ²(3, N = 18,066 detected genes) = 1137.6, p < 0.001, five of which were unannotated RNAs (**Figure 1B; Supplementary Table S2**).

Within groups, males and females shared limited DEG concordance in whole blood except for the MPE group (**Figure 1B; Supplementary Table S5**). In PPE blood, 10 DEGs overlapped between sexes, six (60%) of which were expressed in opposite directions, χ²(1, N = 18,280 detected genes in PPE samples) = 6.5872, p = 0.010 (**Figure 1B; Supplementary Table S2**). MPE males and females shared 229 DEGs, χ²(1, N = 19,620 detected genes) = 1532, p < 2.2*10^-16^, with only 22 (9.6%) expressed in opposite directions (**Figure 1B, Supplementary Table S2**). Of note, *apolipoprotein H* (*Apoh*), and *albumin* (*Alb*) were upregulated in both sexes following MPE, whereas *Adipoq* was upregulated in female, but downregulated in male MPE (**Supplementary Table S2**). Male and female BPE F1s shared 13 DEGs, χ²(1, N = 15,862 detected genes) = 12.9, p = 3.2*10^-4^ (**Figure 1B; Supplementary Table S2**), eight (61.5%) of which were expressed in opposite directions.

Functional enrichment analysis of blood DEGs by Enrichr (referencing the Reactome 2024 database) and Ingenuity Pathway Analysis (IPA) revealed significantly (p_nominal_ < 0.05) enriched pathways that differed across groups, but shared common themes of cell cycle regulation, plasma lipoproteins, DNA repair, immunity, and blood clotting. In PPE males, top terms referenced plasma lipoproteins, cell growth, proliferation, and survival, the extracellular matrix (ECM), immunity, hepatic fibrosis, and blood clotting (**Figure 1C-D; Supplementary Table S3**). PPE females displayed enrichment of vascular endothelial growth factor (VEGF), DNA repair, cell cycle regulation, cancer, Gα_s_ signaling, and ubiquitinol-10 biosynthesis terms (**Figure 1C-D**, **Supplementary Table S3**). Male and female MPE DEGs were enriched for multiple metabolic, platelet-, and blood clotting-related terms, along with plasma lipoproteins, insulin-like growth factor (IGF), acute phase response signaling, and liver X receptor/retinoid X receptor (LXR/RXR) signaling. MPE male DEGs were also enriched for bile acids and salts, whereas MPE females showed additional enrichment for tyrosine catabolism and post-translational protein phosphorylation (**Figure 1C-D**, **Supplementary Table S3**). Across sexes, BPE enrichment included ECM and cell-cell signaling terms. BPE males had additional enrichment for Met, DNA strand elongation, plasma lipoproteins, and collagen. BPE female blood was also enriched for syndecan interactions, muscle contraction, cholesterol biosynthesis, ion homeostasis, and heme degradation (**Figure 1C-D**, Supplementary Table S3).

### mPFC Differential Gene Expression

3’Tag-Seq on F1 mPFC tissue collected from the same mice assessed for whole-blood gene expression revealed 28,697 genes with nonzero counts across groups (**Supplementary Table S4**). A greater proportion of ncRNA and TEC-designated RNA DEGs were represented in mPFC relative to blood; across groups, protein-coding RNAs accounted for ∼50% of DEGs, followed by lncRNAs (∼20-25%), pseudogenes (∼10-15%), TEC-designated RNAs (∼10-15%), and various other ncRNAs (∼1% or less; **Table 2**). DEGs were approximately split between up- and downregulation, apart from greater upregulation of MPE and BPE female mPFC DEGs (∼65%; **Table 2**). Top DEGs by p_nominal_ included genes related to transcriptional regulation, development, transmembrane transport, and synaptic plasticity (**Figure 2A**).

**Figure 2.**
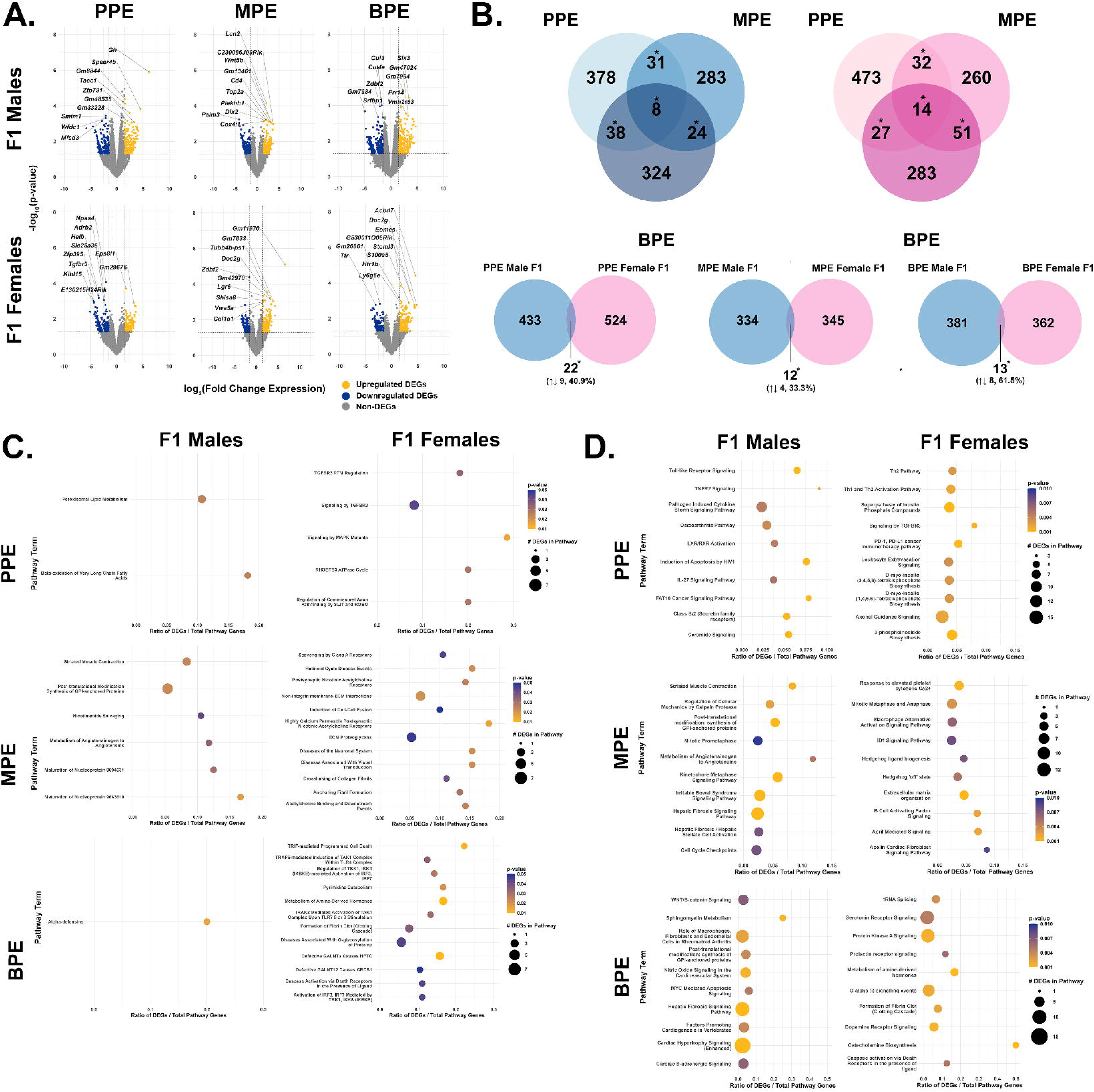
Adult F1 mPFC displays unique differential gene expression signatures based on sex and mode of preconception ethanol exposure. A.) Volcano plots highlighting differentially expressed genes (DEGs), organized by sex and exposure group. DEG thresholds of p < 0.05 and |log_2_FC| > 1.5 are indicated with dotted lines. (B) Venn diagrams comparing DEGs: (top) across exposure groups, within sexes and (bottom) between males and females within exposure groups. All overlaps in gene expression indicated with * were significant by χ² analysis, p < 0.01. (C.) DEG Functional enrichment analysis results from Enrichr utilizing the Reactome 2024 database, organized by sex and exposure group. Top ten terms by p-value are displayed. (D.) DEG functional enrichment analysis results from Ingenuity Pathway Analysis, organized by sex and exposure group. In (C.) and (D.), p-values were truncated at p = 0.001 for ease of display and to ensure a consistent color gradient between males and females.

**Table 2.**
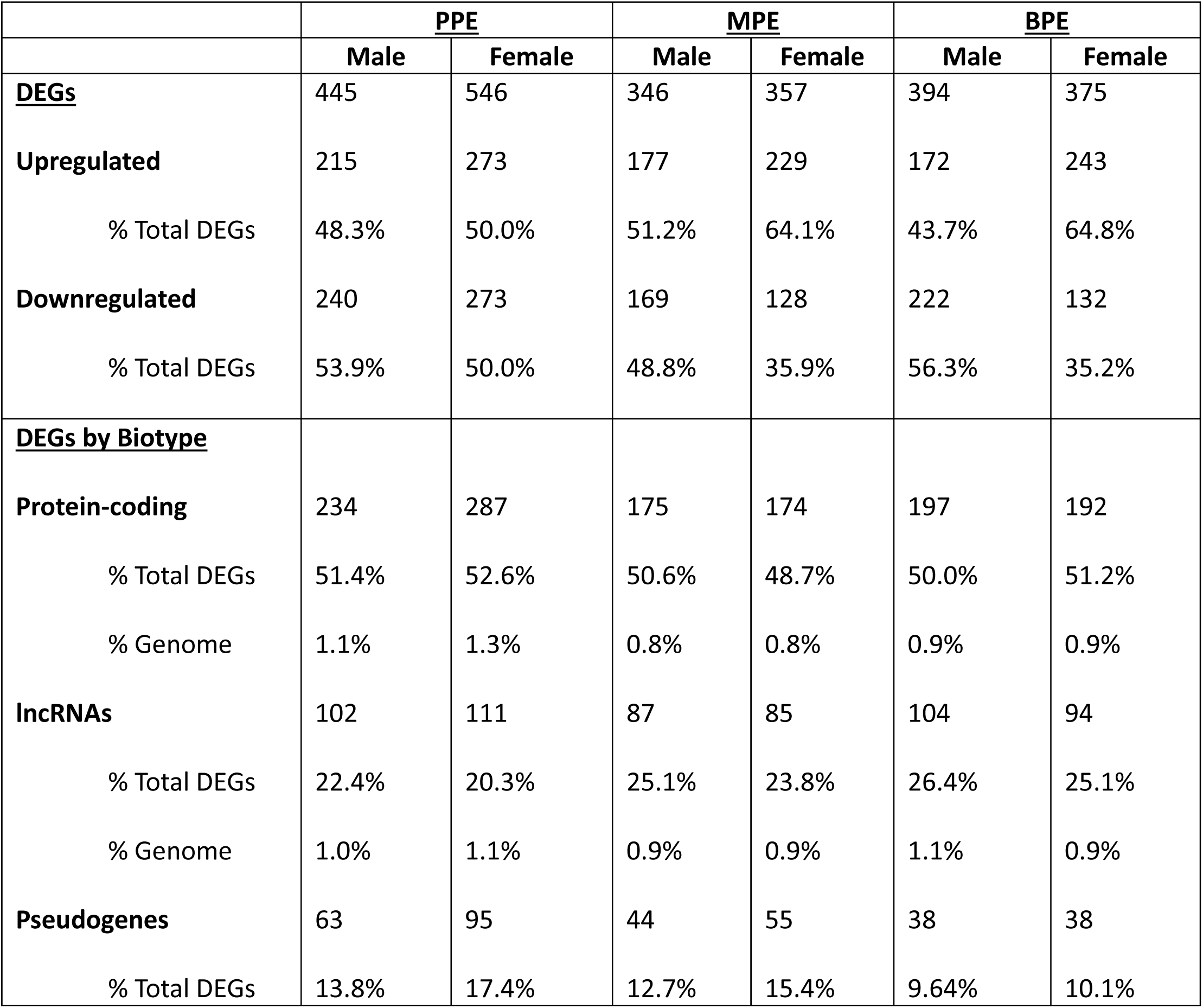
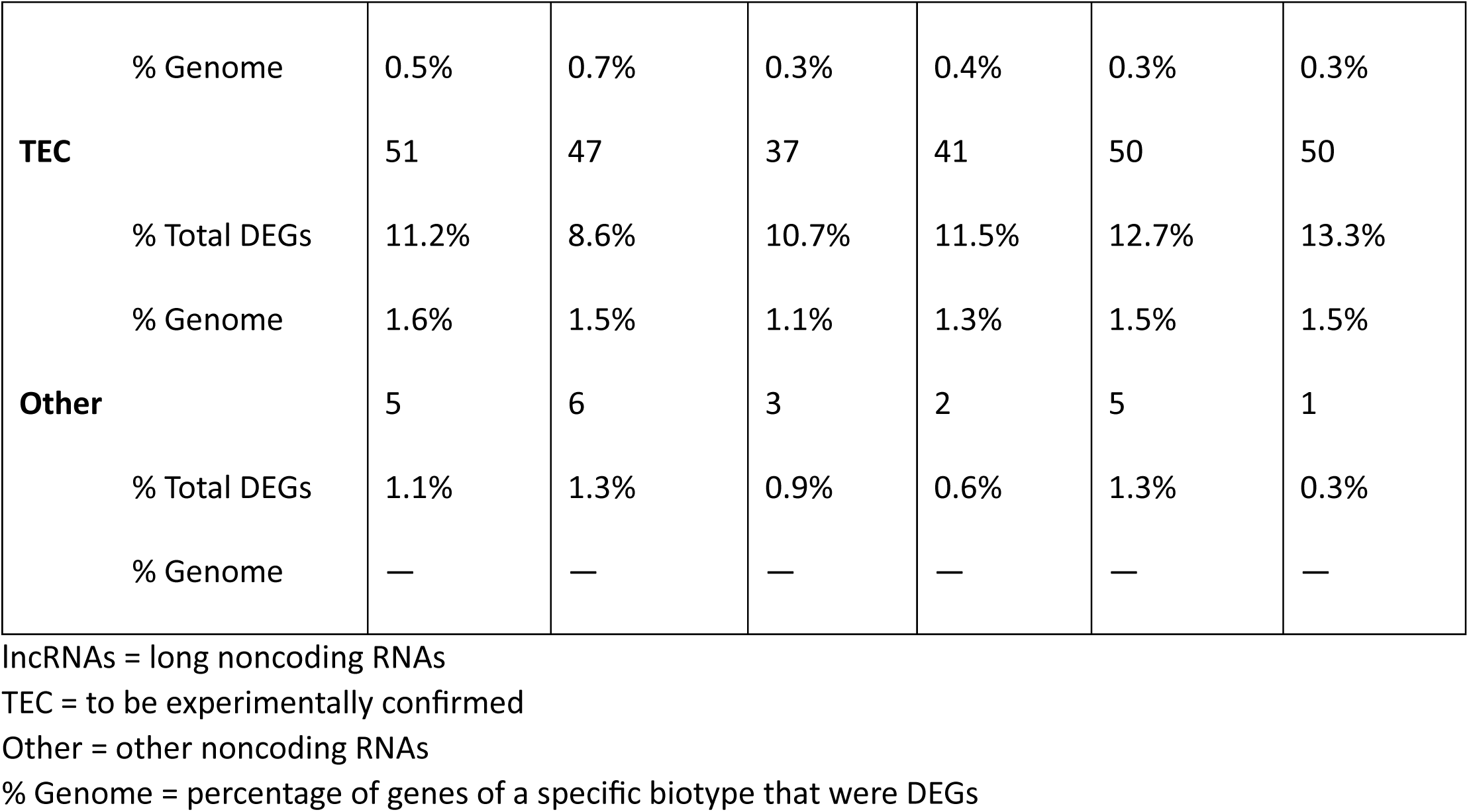
mPFC differential gene expression.

As seen in blood, all overlaps in mPFC DEGs were limited but significant across groups/within sexes and between sexes/within groups. PPE, MPE, and BPE shared only eight DEGs in males and 14 DEGs in females (**Figure 2B; Supplementary Table S4**). PPE males and females shared 22 DEGs, χ²(1, N = 22,292 detected genes) = 10.07, p = 1.5*10^-3^; Nine (40.9%) shared DEGs were expressed in opposite directions (**Figure 2B; Supplementary Table S4**). MPE males and females shared 12 DEGs, χ²(1, N = 25,124 detected genes) = 9.07, p = 2.6*10^-3^, four (33.3%) of which had opposing expression. Thirteen DEGs overlapped between BPE males and females, χ²(1, N = 23,920 detected genes) = 6.69, p = 9.7*10^-3^, eight (61.5%) of which were expressed in opposite directions (**Figure 2B; Supplementary Table S4**).

Functional enrichment of DEGs in mPFC yielded overarching signatures of neuroimmune and neurotransmitter signaling and synaptic regulation, with additional unique pathway terms by group and sex. PPE male DEGs yielded terms related to peroxisomal lipid metabolism and Toll-like receptor (TLR) and tumor necrosis factor (TNF) signaling. In PPE females, enrichment pertained to transforming growth factor beta 3 receptor (Tgbr3) signaling, inositol phosphate biosynthesis, and axon guidance signaling.

Both PPE sexes had enrichment related to inflammatory cytokine signaling (**Figure 2C-D; Supplementary Table S5**). MPE male DEGs were enriched for metabolism of angiotensinogen to angiotensin, post-translational modification synthesis of GPI-anchored proteins, the ECM, cell cycle regulation, and inflammation. MPE enrichment was partially attributable to the differential expression of genes encoding several nicotinic acetylcholine receptor (nAChR) subunits (*Chrna3, Chrna7*) as well as several collagen subunits (*Col1a1, Col4a3*); Other terms related to immunity and cell-growth signaling (**Figure 2C-D**; **Supplementary Table S5**). Top enrichment terms in BPE males were overwhelmingly driven by multiple DEGs encoding voltage-gated calcium channel subunits, the *Wnt* family member *Wnt10b, Tgfbr3,* phospholipase C gamma 2 (*Plcg2*), *nerve growth factor receptor* (*Ngr*), *Myc*, and *Adrb2* (**Figure 2C-D**, **Supplementary Table S5**). BPE females primarily displayed enrichment related to TLR signaling, including terms mentioning TIR-domain-containing adapter-inducing interferon-β (Trif) and interferon regulatory factors 3 and 7 (Irf3, Irf7; **Figure 2B**, **Supplementary Table S5**). Additional enrichment related to catecholamine biosynthesis, serotonin receptor signaling, and Gα_i_ signaling (**Figure 2C**, **Supplementary Table S5**).

### PPE Embryo Differential Gene Expression

To probe the developmental transcriptome of preconception ethanol exposure, we analyzed previously-unpublished male and female PPE embryo RNA-sequencing data obtained during a prior study in our laboratory.^18^ We detected 38,840 genes with nonzero counts across PPE embryos (**Supplementary Table S6**). In both sexes, protein-coding genes accounted for ∼55-65% of DEGs, followed by lncRNAs (∼25%), TEC-designated genes (∼8-12%), pseudogenes (∼8-9%), and various other ncRNAs (∼2%; **Table 3; Supplementary Table S6**). DEGs were modestly downregulated in both sexes (**Table 3**; **Figure 3A**). Top DEGs by p_nominal_ included several unannotated RNAs along with genes encoding transcription factors, ion channels, receptors, and cell-adhesion molecules (**Figure 3A**). Male and female PPE embryos shared 95 DEGs, χ²(1, N = 38,840 detected genes) = 179.5, p < 2.2*10^-16^, 40 (42.1%) of which had opposing expression (**Figure 3B; Supplementary Table S6**). Shared DEGs of note included *C-X-C motif chemokine receptor 5* (*Cxcr5,* upregulated in male, downregulated in female embryos), *leukocyte receptor tyrosine kinase* (*Ltk,* downregulated in both sexes), and *transcription factor AP-2 beta* (*Tfap2b*, upregulated in both sexes; **Supplementary Table S6**).

**Figure 3.**
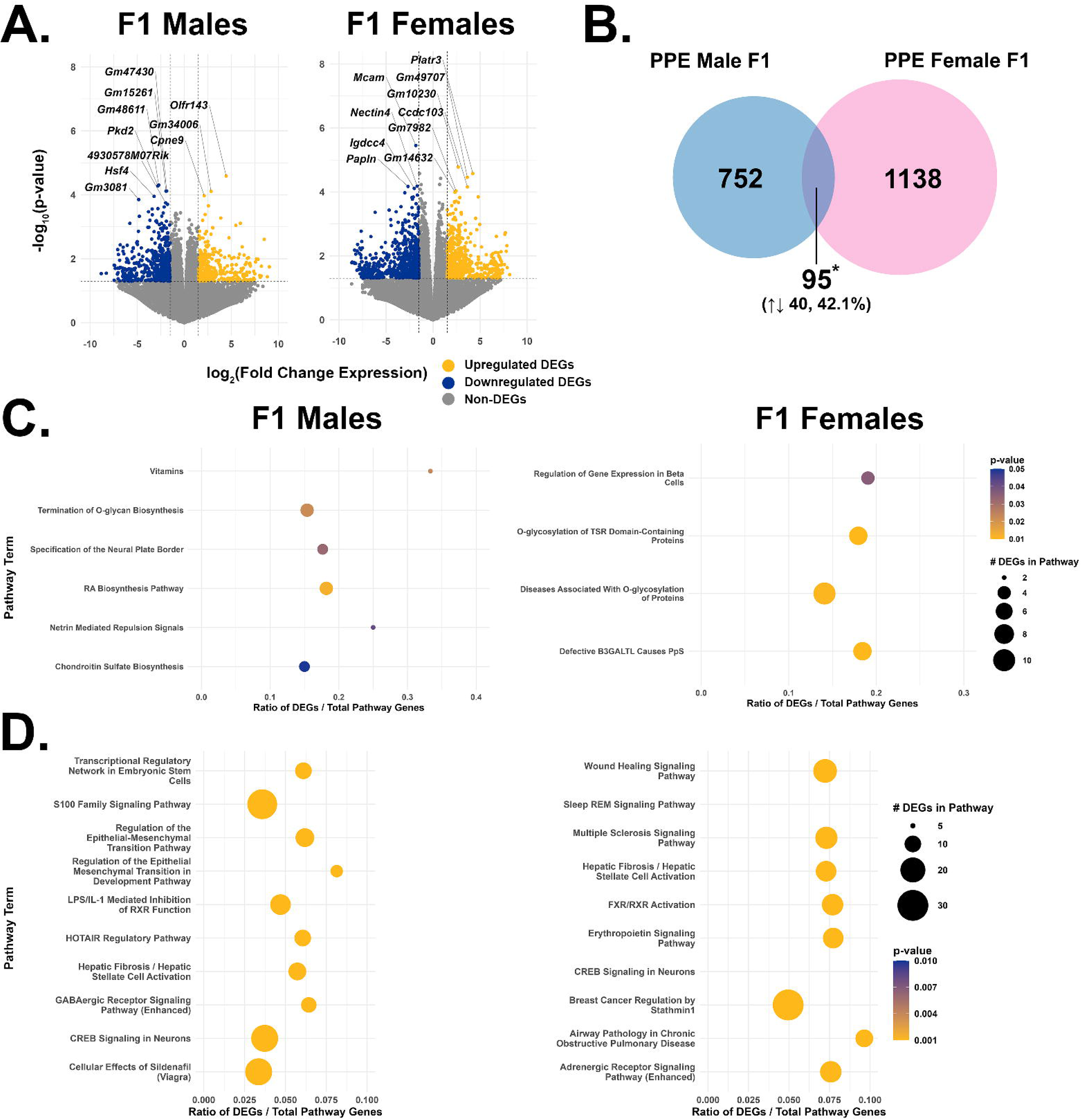
F1 PPE embryos show sex-specific patterns of gene expression. (A.) Volcano plots highlighting differentially expressed genes (DEGs), organized by sex. DEG thresholds of p < 0.05 and |log_2_FC| > 1.5 are indicated with dotted lines. (B) Venn diagram comparing DEGs between males and females within exposure groups. The overlap in DEGs between male and female PPE embryos was significant by χ² analysis, p < 2.2*10^-16^. (C.) DEG Functional enrichment analysis results from Enrichr utilizing the Reactome 2024 database, organized by sex. Top ten terms by p-value are displayed. (D.) DEG functional enrichment analysis results from Ingenuity Pathway Analysis, organized by sex. In (C.) and (D.), p-values were truncated at p = 0.001 for ease of display and to ensure a consistent color gradient between males and females.

**Table 3.** Embryo differential gene expression.

|  | **PPE** |  |
| --- | --- | --- |
|  | **Male** | **Female** |
| **<u>DEGs</u>** | 847 | 1233 |
| **Upregulated** | 305 | 537 |
| % Total DEGs | 36.0% | 43.5% |
| **Downregulated** | 542 | 696 |
| % Total DEGs | 63.9% | 56.4% |
| **<u>DEGs by Biotype</u>** |  |  |
| **Protein-coding** | 435 | 708 |
| % Total DEGs | 51.4% | 57.4% |
| % Genome | 1.9% | 3.2% |
| **lncRNAs** | 212 | 296 |
| % Total DEGs | 25.0% | 24.0% |
| % Genome | 2.1% | 3.0% |
| **Pseudogenes** | 80 | 100 |
| % Total DEGs | 9.4% | 8.1% |
| % Genome | 0.6% | 0.7% |
| **TEC** | 102 | 106 |
| % Total DEGs | 12.0% | 8.6% |
| % Genome | 3.2% | 3.3% |
| **Other** | 18 | 23 |
| % Total DEGs | 2.1% | 1.8% |
| % Genome | — | — |

lncRNAs = long noncoding RNAs
TEC = to be experimentally confirmed
Other = other noncoding RNAs
% Genome = percentage of genes of a specific biotype that were DEGs

Functional enrichment results in PPE embryos were broadly attributable to DEGs encoding components of various signaling pathways, including multiple G protein-coupled receptors and their ligands, cytoskeletal components, protein kinases, and cytokines and their receptors (**Figure 3C-D**, **Supplementary Table S7**). Notably, “specification of the neural plate border” was enriched in PPE male embryos, driven by *transcription factor AP-2 beta* (*Tfap2b*)*, transcription factor 7 like 1* (*Tcf7l1*), and *gastrulation brain homeobox 2* (*Gbx2*; **Figure 3C**, **Supplementary Table S7**). Three of the four enriched terms in female PPE embryos (“Defective B3GALTL Causes PpS”, “O-glycosylation of TSR Domain-Containing Proteins”, and “Diseases Associated With O-glycosylation of Proteins”) were driven by DEGs in the ADAMTS family.

### Integrative Analysis of Differential Expression Results

Expanded comparisons of DEGs across sexes and tissues did not reveal meaningful overlaps. No DEGs were shared across all group/sex/tissue combinations, within sex across groups/tissues, nor within sex/group across tissues (**Figure 4A**). Although individual DEGs may not overlap across sexes and/or tissues, the same biological pathways could be affected through different DEGs. As such, we also compared enriched (p_nominal_ < 0.05) pathways from gene set enrichment analysis (GSEA) referencing all 238 Enrichr databases across all group/sex/tissue combinations. While this approach did not improve overlaps across groups/sexes/tissues, it yielded overlapping pathways within sex/group combinations across blood and mPFC (**Figure 4B**). We combined genes and GSEA statistics from shared pathways across blood and mPFC within group/sex combinations (**Supplementary Table S8**) and found several notable shared pathways. In PPE males, the top shared blood-mPFC pathways by average p_nominal_ related to preclinical models of Huntington’s disease from the Huntington’s Disease in High Definition database (HDSigDB; **Supplementary Table S8**). Additional HDSigDB terms were enriched in this dataset and in shared PPE female tissue pathways. The top enriched shared pathway in MPE males was from the GO Cellular Component 2015 database and related to the extracellular region (**Supplementary Table S15**). MPE female tissues shared many pathway terms related to liver and small intestine, driven by plasma lipoproteins and metabolic genes (*e.g. Fabp1, Apoa1, Aldob, Angptl3*) in blood and by cell-growth and immune-related genes (*e.g., Cenpii, Kntc1, Tnfrsf17*, *Il12rb1*) in mPFC (**Supplementary Table S8**). BPE male tissues shared enriched pathway terms from multiple chromatin immunoprecipitation-sequencing databases, denoting changes in epigenetic regulation of gene expression, including several terms pertaining to the histone acetyltransferase P300 along with the RNA polymerase II subunit Pol2 and the transcription factor Stat3 (**Supplementary Table S8**). Like PPE males and females, BPE female tissues shared multiple enriched terms related to models of Huntington’s disease (**Supplementary Table S8**).

**Figure 4.**
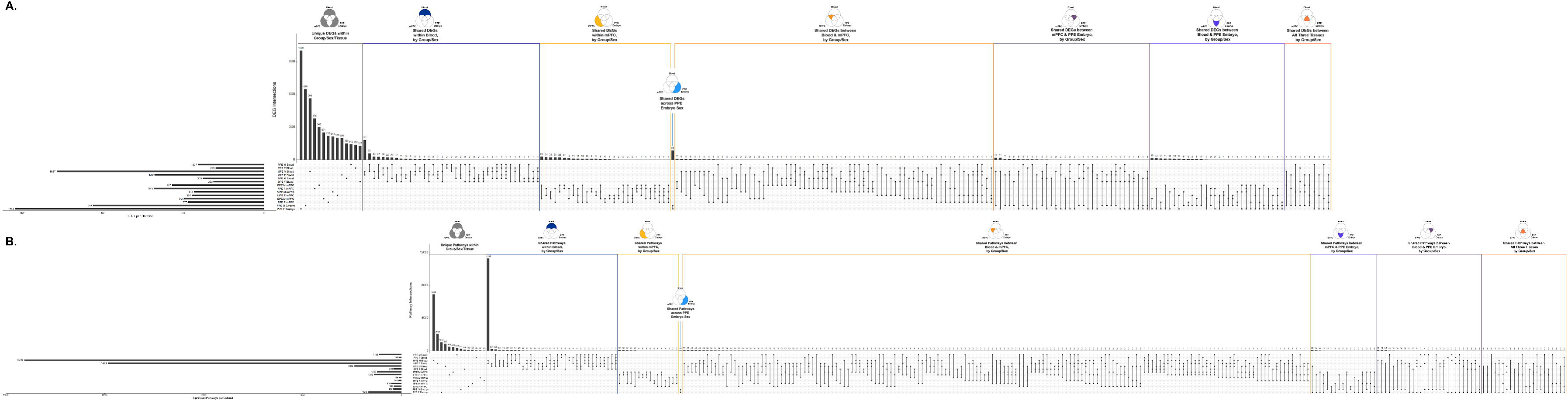
DEGs and pathways poorly intersect across sexes, exposures, and tissues. UpSet plots displaying intersections across datasets (encompassing all group/sex/tissue combinations) of (A.) DEGs and (B.) Pathways from gene set enrichment analysis referencing all 238 databases from Enrichr. Horizontal bar plots on the left side depict the total number of genes or pathways associated with each dataset. Individual dots in the main region of the plot are unique genes or pathways to their respective dataset. Each set of dots connected by vertical lines can be understood as a unique intersection within a Venn diagram comparing the associated datasets on the y-axis, *i.e.,* the overlapping elements in this intersection (DEGs or pathways) are not present in any other intersection. The vertical barplot depicts the total number of genes or pathways in each comparison. From left to right, the first region shows unique genes or pathways, the second region shows genes or pathways only shared between blood datasets, the third shows genes or pathways only shared between mPFC datasets, the fourth shows genes/pathways uniquely shared between male and female PPE embryos, the fifth shows genes/pathways shared between blood and mPFC datasets, the sixth shows genes/pathways shared between mPFC and PPE embryo datasets, the seventh shows genes/pathways shared between blood and PPE embryo datasets, and the final shows DEGs/pathways between all three tissues’ datasets.

### Rank-Rank Hypergeometric Overlap (RRHO2)

We next utilized the RRHO2 algorithm to compare transcriptome-wide gene expression between sexes, within tissues and groups.^20^ Except for the PPE embryo and MPE blood groups, all comparisons had -log_10_(p-value) ranges <100, most of which were <20 (**Figure 5A-C**), indicating weak overlapping gene expression signatures between datasets.^20^ MPE whole blood comparisons reached a peak -log_10_(p-value) of 178, with most genes falling around -log_10_(p-value) <36 (**Figure 5B**). PPE male and female embryos showed the greatest level of overlapping gene expression, with majority concordant gene expression reaching greater log_10_(p-value) ranging ∼150-392 (**Figure 5D**).

**Figure 5.**
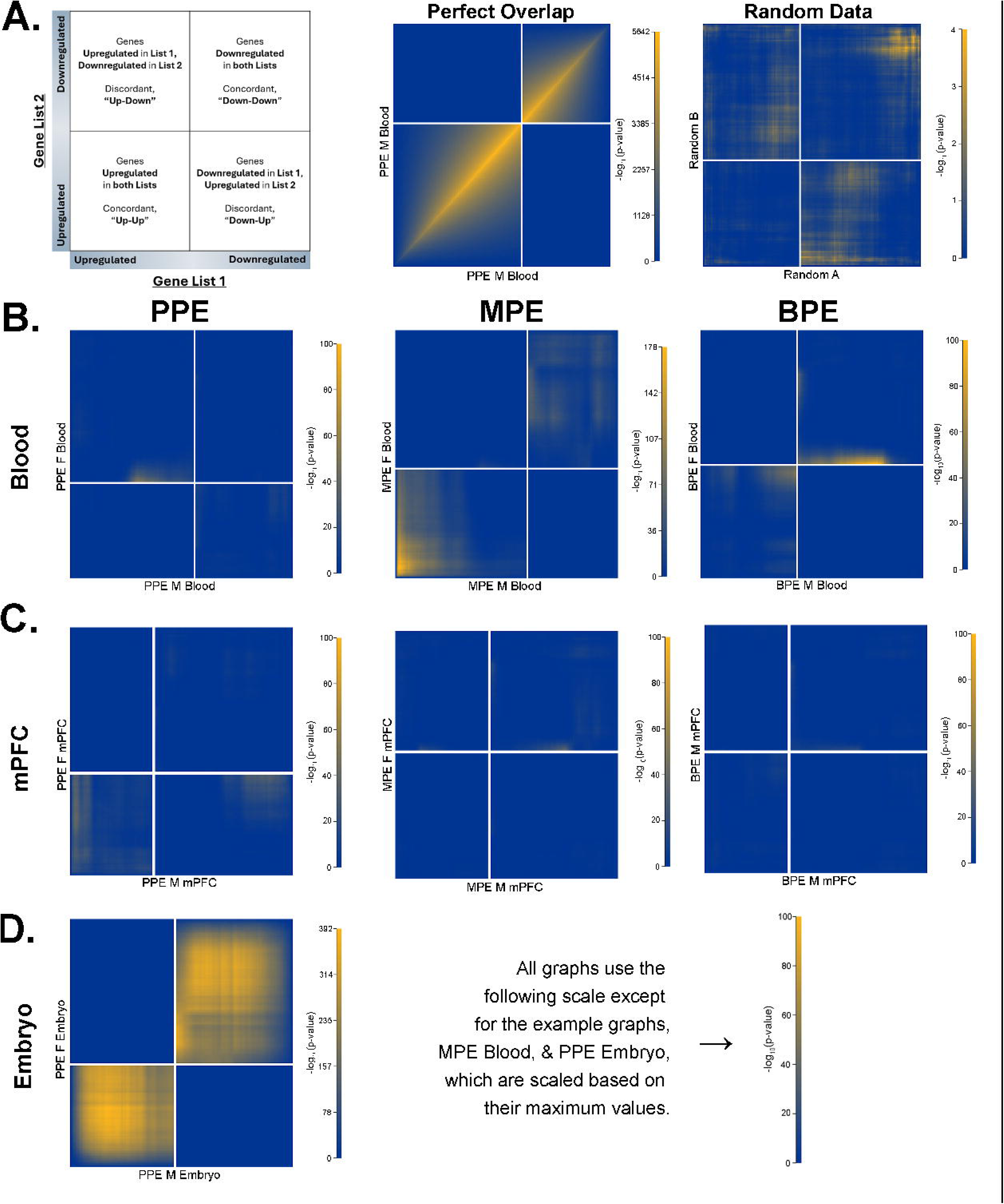
Robust sex differences in F1 transcriptomes persist within groups and tissues using a threshold-free approach (RRHO2). (A.) Legend. The leftmost image demonstrates how to interpret the plots’ four quadrants. Subsequent plots demonstrate a perfect overlap (middle, comparing the PPE male offspring whole blood dataset to itself) and simulated random data (based on the minimum and maximum raw counts of PPE male offspring whole blood data). (B.) RRHO2 graphs comparing male and female expression data from whole blood. (C.) RRHO2 graphs comparing male and female expression data from mPFC. (D.) RRHO2 graph comparing male and female PPE embryo data.

### Correlation of Blood & mPFC Gene Expression

To determine whether offspring whole blood could predict mPFC gene expression, we performed within- and between-subjects correlations across tissues/within sexes, following the methods of Ferguson *et al.* (2022).^21^ As they examined the F0 generation,^21^ we compared genes with significant within-subjects blood-mPFC correlations from their study to those identified here to pinpoint potential biomarkers of chronic ethanol exposure persistent across generations. Due to our relatively small sample sizes (n = 5-7 samples/sex/group), for generalizability of correlations across parental exposures, and for comparability with Ferguson *et al.*,^21^ between- and within-subjects correlations were collapsed across exposure groups/within sexes.

Between-subjects correlations revealed a significant (Holm-Bonferroni p_adj_ < 0.001) correlation of modest strength (males, Spearman ρ = 0.35; females, ρ = 0.36) between whole blood and mPFC gene expression in both sexes (**Figure 6A**), suggesting a limited, but present correspondence of gene expression between tissues. Within-subjects correlations yielded 607 genes with significant (p_adj_ < 0.05) blood-mPFC correlations for males and 653 genes for females (**Supplementary Table S9**). Of these genes, 46 (7.58%) were also whole-blood DEGs in at least one exposure group in males, and 27 (4.13%) in females (**Supplementary Table S9**). The top correlated genes also differentially expressed in blood were *TBC1 domain family member 32* for males (*Tbc1d32;* Spearman ρ = 0.66, p_adj_ < 0.001) and *Gm8744* for females (Spearman ρ = -0.59, p_adj_ < 0.01; **Supplementary Table S9**).

**Figure 6.**
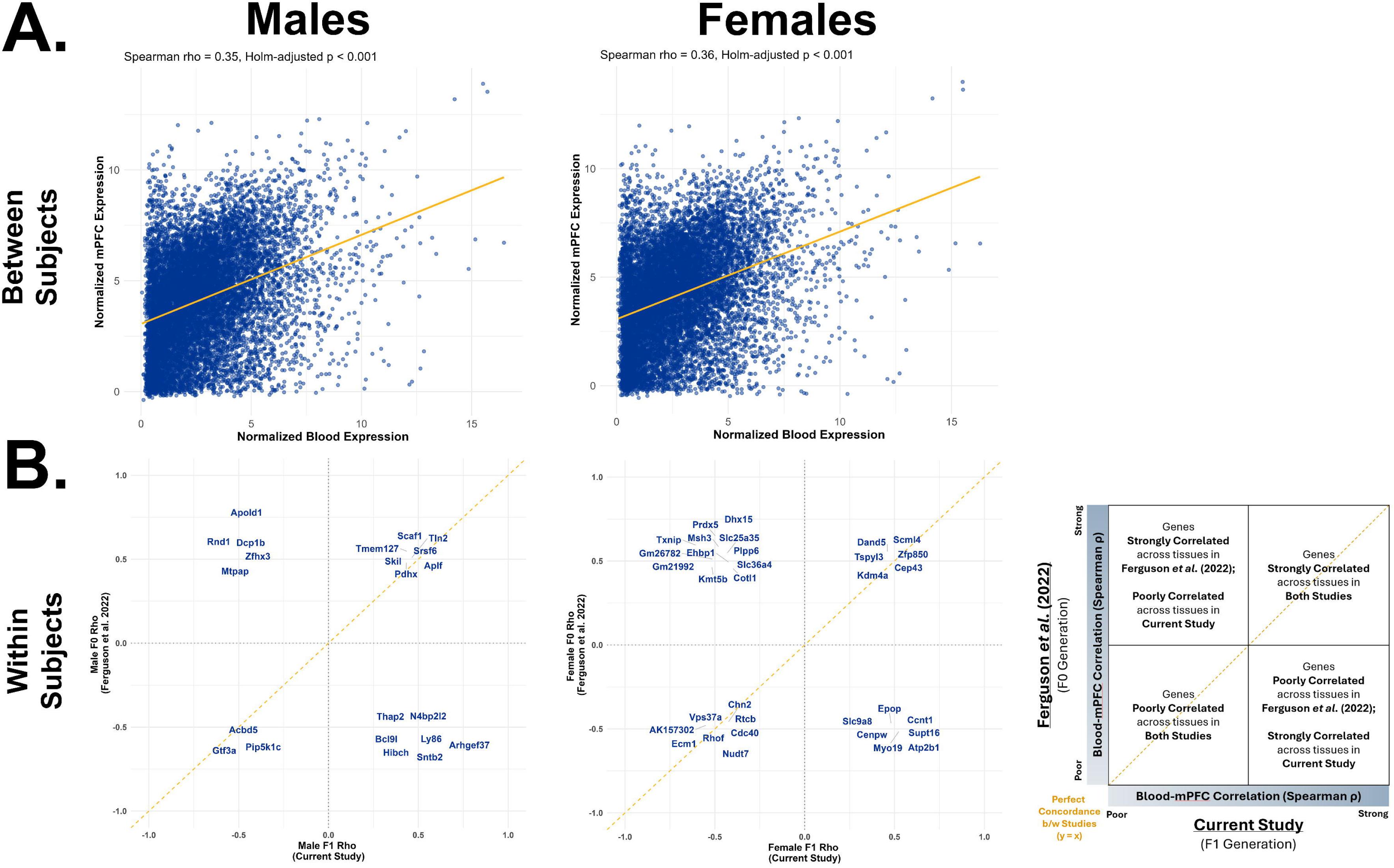
Whole blood and mPFC transcriptomes weakly correlate in F1 with preconception ethanol exposure. (A.) Between-subjects correlations for normalized expression of individual genes between whole blood and mPFC, averaged across subjects in (left)males and (right) females. (B.) Cross-study comparison of significant (Holm-Bonferroni p_adj_ < 0.05) within-subjects correlations of normalized gene expression between whole blood and mPFC within (left) males and (right) females between the current study and Ferguson *et al.* (2022).

Comparing within-subjects blood-mPFC correlations between F1 mice in the current study to those in F0 from Ferguson *et al.*^21^ revealed a limited overlap in gene expression signatures. There were 22 genes with significant within-subjects blood-mPFC correlations across studies for males (3.62% of our total significant male within-subjects correlations) and 33 overlapping genes for females (5.20% of significant female within-subjects correlations; **Figure 6B**, **Supplementary Table S9**). For males, 10 (45.5%) of these genes were concordant across studies, whereas 12 (57.14%) were discordant (**Figure 6B**, **Supplementary Table S9**). For females, 14 genes (42.4%) were concordant while 19 (57.6%) were discordant (**Figure 6B**, **Supplementary Table S9**). In males, only *talin 2* (*Tln2*) was shared between studies and was also a whole-blood DEG in at least one group in the current study (**Figure 6B**, **Supplementary Table S9**). In females, three shared genes across studies were also whole-blood DEGs: *centromere protein W* (*Cenpw*), *nudix hydrolase 7* (*Nudt7*), and *solute carrier family 25 member 35* (*Slc25a35*; **Figure 6B**, **Supplementary Table S9**).

### Machine Learning Classification of Preconception Ethanol Exposure from Whole-Blood Gene Expression

Finally, we sought to establish whether whole-blood gene expression could predict preconception ethanol exposure status using machine learning classification algorithms. This also followed the methods of Ferguson *et al.*^21^ We collapsed subjects within sexes/across groups to predict overall preconception ethanol exposure status without overfitting to a single mode of exposure. We assessed model performance via the area under the receiver operating characteristic curve (AUC), comparing random forest (RF), logistic regression with elastic net regularization (LR), partial least squares discriminant analysis (PLSDA), and extreme gradient boosting (XGB) models. In male offspring, whole blood was a weak predictor of preconception ethanol exposure status, with most models performing marginally better than random chance (AUC = 50%): RF yielded AUC 64.2%, LR 58.8%, PLSDA 50.7%, and XGB 59.5% (**Figure 7A**). Models had moderate performance in female offspring whole blood, with RF producing AUC 78.8%, LR 65.3%, PLSDA 67.5%, and XGB 62.9% (**Figure 7A**).

**Figure 7.**
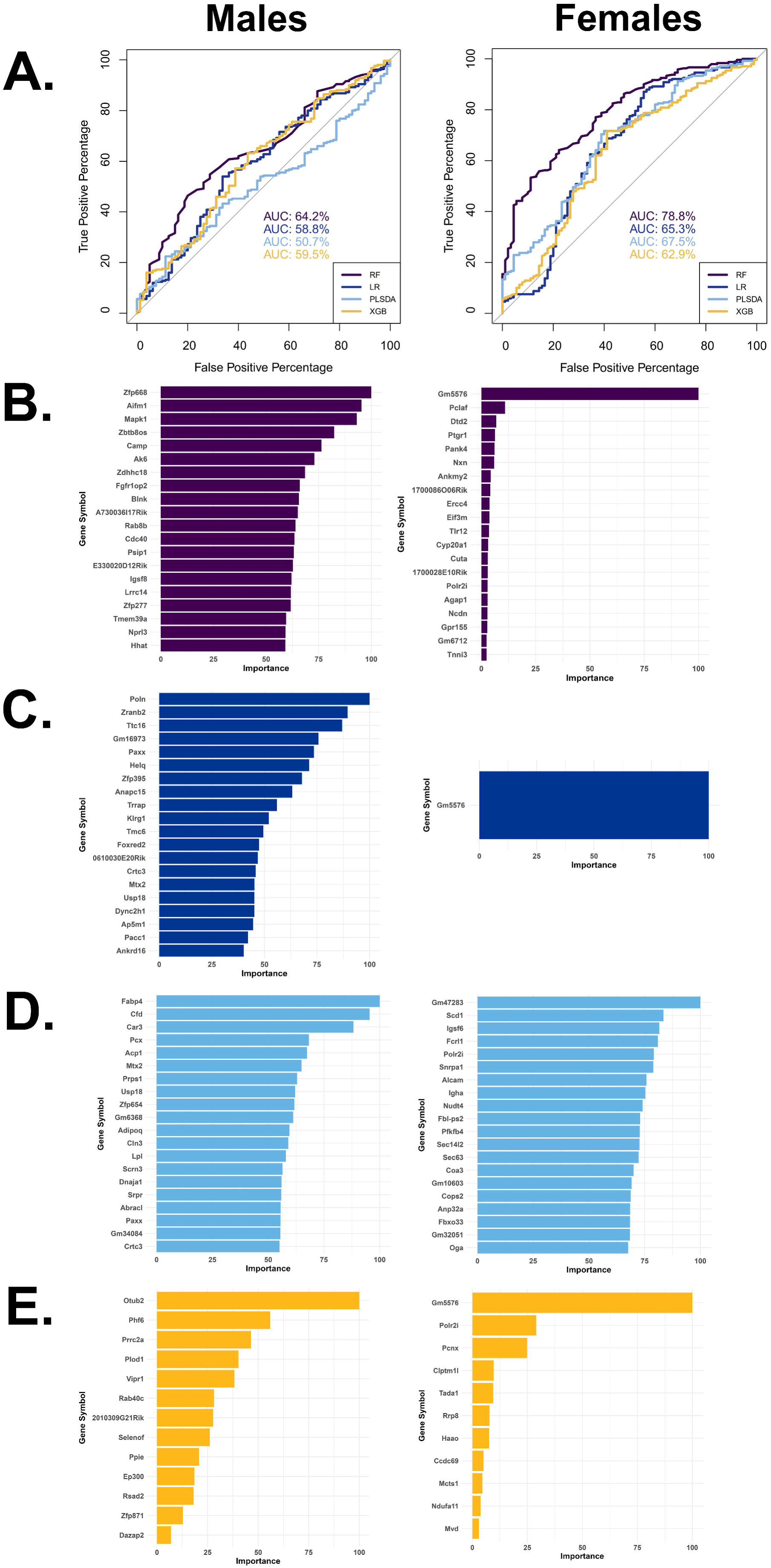
The F1 whole blood transcriptome is a poor predictor of preconception ethanol exposure status across machine learning classifiers. (A.) Receiver operating curves and AUC values for (left) males and (right) females using random forest (RF) and logistic regression with elastic net regularization (LR), partial least squares discriminant analysis (PLSDA) within the MLSeq R package as well as the extreme gradient boosting (XGB) model from the caret R package. (B.-D.) Top 20 most important genes for preconception ethanol exposure status classification for (B.) RF, (C.) LR, (D.) PLSDA, (E.) XGB models.

The top 20 most important genes selected by each model during training are presented in **Figure 7B-E**; All genes with nonzero variable importance are listed within **Supplementary Table S10**. No gene overlapped across all models’ nonzero features for males, but the unannotated noncoding gene *Gm5576* overlapped across all models for females with high importance and was the single gene utilized by the LR model to classify preconception ethanol exposure status (**Figure 7B-D**, **Supplementary Table S10**).

*Gm5576* was a shared whole-blood DEG downregulated across groups in females (**Supplementary Table S4**), but it was not found to correlate in expression between blood and mPFC in within-subjects correlations (**Supplementary Table S10**). Of note for males, the RF model, which had the greatest performance, selected the unannotated noncoding gene *E330020D12Rik* to determine preconception ethanol exposure status, which was a DEG shared between PPE, MPE, and BPE male whole blood (**Figure 7B**, **Supplementary Table S3**, **Supplementary Table S10**).

## Discussion

The current study comprehensively probed the transcriptomes of ethanol-naïve F1 offspring with chronic preconception ethanol exposure, comparing RNA-sequencing data across male and female PPE, MPE, and BPE whole blood and mPFC and synthesizing these findings with PPE embryo data obtained during our prior study.^18^ Based on available literature, we expected some group- and sex differences in the transcriptomic effects of preconception ethanol, though we predicted commonalities in affected pathways. Our data revealed highly unique signatures of preconception ethanol exposure based on parental exposure, offspring sex, and tissue analyzed. The sex-dependent nature of cross-generational transcriptomic outcomes of chronic preconception ethanol exposure alone constitutes a valuable conclusion from the current study. Although specific DEGs and pathways diverged across exposure groups and sex, shared themes emerged from our analyses.

PPE-embryo transcriptomic data revealed the effects of chronic preconception ethanol emerge within the first developmental days. Many DEGs were transcription factors, including *Tfap2b, Tcf7l1,* and *Gbx2*, which play critical roles in craniofacial, central nervous system, and cardiovascular development.^22–25^ Other DEGs included chemokines and their receptors, further indicating altered developmental signaling pathways; *Cxcr5* and *Ltk,* implicated in neuronal migration and polarization,^26, 27^ were shared between sexes. These data support findings from other PPE studies, which noted similar offspring phenotypes to those characteristic of fetal alcohol spectrum disorders (FASDs), including altered developmental programming,^12^ growth restriction,^11, 28^ craniofacial growth abnormalities,^13^ and aberrant nervous system development.^29^

Though altered developmental gene expression may explain some adult phenotypes of preconception ethanol exposure, this must be validated with causative studies. Our findings present candidate genes and pathways to investigate in future research. For example, *Met* was differentially expressed in PPE embryos in opposite directions by sex, while terms related to Met signaling displayed functional enrichment in PPE and BPE male-, and MPE male and female blood. *Met* encodes a hepatocyte growth factor-activated tyrosine receptor kinase, and is implicated in the development of multiple cancers and in diabetes.^30, 31^ Indeed, gene expression patterns related to metabolic dysfunction were observed across groups and sexes; many of these DEGs play additional roles in immunity and inflammation. *Cfd, Cidec, Adipoq, Fabp4, Apoh,* and their respective protein products are secreted by adipocytes or hepatocytes into the bloodstream, which can contribute to the pathogenesis of metabolic and cardiovascular diseases and certain cancers.^32–36^

Prior studies have noted sex-specific metabolic dysfunction in offspring with preconception ethanol exposure. Male-specific PPE effects have been found on growth restriction during fetal and early postnatal development, which were associated with adult insulin hypersensitivity and upregulation of pro-fibrotic- (including genes in the TGF-β signaling pathway) as well as LXR/RXR signaling pathway genes in liver.^28^ Another study found elevated liver LXR signaling modulates resistance to obesity, along with improved glucose homeostasis following a high-fat diet challenge.^37^ We found LXR/RXR signaling pathway enrichment across offspring sex and parental-exposure groups, corroborating results in other modes of preconception ethanol exposure. Human studies consistently link heavy alcohol consumption and AUD with increased risk of type II diabetes^38, 39^ and alcohol-associated liver disease (ARLD).^40^ Both diseases have (epi)genetic components,^41, 42^ but human studies specifically examining their heritability with familial AUD are limited. Although AUD is associated with impaired metabolic phenotypes, familial studies have focused on alcohol metabolism, namely transmission of gene variants encoding the alcohol-and acetaldehyde dehydrogenase enzymes in AUD and ARLD risk.^41^

The abundance of whole-blood DEGs expressed by liver and adipocytes is notable, but not unexpected. Organs secrete RNA and protein into the blood via extracellular vesicles, which mediate cross-tissue communication and exhibit altered cargo and release mechanics in pathological states,^43^ such as ARLD, where exosomes and their cargo show promise as diagnostic and therapeutic targets.^44^ Extracellular vesicles can cross the blood-brain barrier and facilitate bidirectional communication between the brain and other organs.^45, 46^ Thus, whole-blood DEGs with primary expression in non-blood-derived cell types may reflect extracellular vesicle-mediated cross-generational risk or protective phenotypes of alcohol-related disease, though this requires further study. Future studies may isolate specific blood fractions or cell types to analyze their individual gene expression profiles, as has been performed in human and animal FASD research, as well as in human AUD heritability studies.^47–49^ This approach may help eliminate transcriptomic influences of non-blood-derived cell types.

Analyses of the mPFC in PPE, MPE, and BPE offspring revealed transcriptomic effects related to immunity and neurotransmitter signaling across groups and sexes. Immune-related DEGs and pathways represented TLRs, TGF-β, TNF, and inflammatory cytokine signaling. Neuroimmune dysfunction contributes to the pathogenesis and maintenance of AUD,^50^ though to our knowledge no existing studies of preconception ethanol exposure have specifically investigated intergenerational neuroimmune impacts. We also observed differential expression in mPFC of several genes encoding enzymes for monoamine synthesis along with monoamine and acetylcholine receptors. Rodent models of AUD display altered mPFC levels of serotonin,^51^ dopamine,^51^ and norepinephrine.^52^ Altered serotonergic and dopaminergic functioning have been corroborated in frontal cortex of humans with AUD^53, 54^ and are positively correlated with AUD family history.^53, 55^ Early PPE studies found altered levels of norepinephrine, serotonin, and Met-enkephalin in offspring brain.^56^ A more recent prenatal ethanol exposure study additionally found altered nAChR signaling in mPFC.^57^ Our data suggest alterations in mPFC monoamine and acetylcholine transmission may extend to other modes of preconception ethanol exposure in a sex-dependent manner.

Our gene expression comparisons across groups, sexes, and tissues reinforce context-dependent molecular biology of chronic ethanol exposure. DEG comparisons and RRHO2 found limited gene expression concordance between sexes, within exposure groups/tissues. Between-subjects correlations revealed the whole-blood transcriptome is a modest predictor of the mPFC transcriptome in offspring with chronic preconception ethanol exposure, and within-subjects comparisons identified hundreds of individual genes within sexes with moderate-to-strong correlations between tissues. Machine learning classifiers, particularly the RF algorithm, displayed moderate performance in predicting preconception ethanol exposure status in female F1, indicating a moderate, sex-specific persistence of a of a chronic ethanol exposure blood signature. Ferguson *et al.* found moderate-to-strong prediction of CIEV-status based on whole-blood gene expression in F0, with stronger predictive signatures in females and with the LR and PLSDA algorithms.^21^ Although CIEV produces robust gene expression changes in blood and brain, many transcriptomic responses are time- and cell type-specific.^21, 58, 59^ In our cross-study comparison of within-subjects correlations, we identified a limited set of genes in both sexes with correlated blood–mPFC expression in the Ferguson *et al.* F0 dataset^21^ and our F1 dataset. Further studies should compare blood and brain gene expression in ethanol-exposed parents and their offspring in tandem to verify generation-dependent effects of CIEV across tissues, though our analyses uncovered the unannotated ncRNAs *Gm5576* (females) and *E330020D12Rik* (males) as potential targets for future research.

Although no DEGs nor pathways overlapped across all datasets within the current study, various growth factors were notably differentially expressed across groups, sexes, and tissues. VEGF signaling was enriched in PPE female, MPE male and female, and BPE male whole blood. In AUD, VEGF is a biomarker of neuroinflammation and frontal lobe impairment,^60^ and its expression is induced in peripheral blood mononuclear cells of nonhuman primate models of chronic ethanol consumption.^61^ *Gh* was a top upregulated DEG in PPE male and BPE female mPFC, and was upregulated in female PPE embryos along with *growth hormone releasing hormone receptor*, which regulates *Gh* expression. Gh has neurotrophic, neuroprotective, and synaptogenic roles in adult brain,^62^ though a study in male mice found elevated prefrontal cortical *Gh* expression in a diabetes model correlates with poor learning and memory.^63^ We also found differential expression of genes encoding members of the IGF signaling cascade in blood of MPE males and females and BPE males, as well as in BPE female and MPE male mPFC. IGF signaling is involved in ARLD^64^ and FASDs.^65^ Rodent studies have found reduced IGF1 in plasma and in multiple brain regions following chronic ethanol exposure, effects also associated with neuron loss.^66, 67^ IGF signaling is also associated with neuronal loss in multiple brain regions^67^ and cognitive impairments^68^ in individuals with AUD.

After protein-coding genes, lncRNAs were the next most abundant biotype across F1 groups and tissues. Many of these lncRNAs remain functionally uncharacterized but display marked differential expression with preconception ethanol exposure and/or were shared across groups or sexes. Prior research links ncRNAs, including lncRNAs, to ethanol-related phenotypes, with robust expression changes in AUD subjects.^69^ Mounting evidence also supports a role for germline ncRNAs in programming preconception ethanol exposure phenotypes.^4^ Some studies have linked differential ncRNA expression in F0 germ cells to offspring gene expression and molecular and behavioral phenotypes, though this has been more extensively investigated in preconception stress and trauma.^4^ Characterizing the noncoding genome will elucidate cross-generational disease risk following preconception ethanol exposure, though additional studies are needed to causally test the role of ncRNAs in these phenotypes.

Our study has several limitations. First, the method and duration of ethanol exposure and the exposure-to-breeding interval may influence offspring phenotypes.^4^ No consensus exists among preconception ethanol studies on exposure paradigms,^4^ though we chose CIEV-2BC for its reliable induction of ethanol dependence-related molecular and behavioral phenotypes^19^ and their translatability across species.^70^ Due to differing CIEV and 2BC cycle lengths (four versus five days), the interval between exposure cessation and breeding differed between sires and dams (72- versus 48 hours, respectively), but remained consistent across groups. Exposures immediately prior to breeding (CIEV in males versus 2BC in females) may affect F1 phenotypes, however, alternating sexes between CIEV and 2BC minimized confounding effects of male-female proximity in vapor chambers. Although litter size and sex ratios were unchanged with preconception ethanol exposure, pup mortality was significantly higher in MPE and modestly, though not significantly, elevated in BPE litters. Once pregnancies were confirmed, sires were removed and litters were monitored during the first three postnatal days. We found pup mortality in the MPE and BPE groups was attributable to maternal infanticide, likely indicative of maternal aggression due to increased stress from CIEV-2BC.^4^ Without utilizing embryo transfer and/or cross-fostering, however, it is difficult to distinguish the phenotypic outcomes in offspring of parental ethanol exposure from those induced by ethanol-induced changes in maternal care or maternal aggression.^4^ Despite these limitations, our analyses revealed novel transcriptomic signatures of PPE, MPE, and BPE in both offspring sexes that can be validated with further preconception ethanol studies.

## Acknowledgements

This project utilized the services of the University of Pittsburgh Health Sciences Sequencing Core at UPMC Children’s Hospital of Pittsburgh for TapeStation RNA fragmentation analysis. RNA Sequencing of whole blood and mPFC samples was performed by the Genomic Sequencing and Analysis Facility at UT Austin, Center for Biomedical Research Support, RRID#: SCR_021713. We acknowledge the award “Cross-generational effects of pre-conception alcohol use and cancer” from Bridging Connections in Addiction and the Pittsburgh Foundation for primary support of this study. We also acknowledge ancillary support from NIAAA grants U01 AA020889 (MPIs: Farris and Homanics), R01 AA030257 (PI: Farris), F31 AA031168 (PI: Rice), and F31 AA032172 (PI: Gil).

## Data Availability

The transcriptomic data underlying this article are available in GEO under accession number GSE329570. All code used for bioinformatic analyses is available upon request.

## Ethics Statement

All animal studies were approved by the Institutional Animal Care and Use Committee at the University of Pittsburgh (approved protocol #23083423).

