## Supplementary Figures for "Preconception Chronic Intermittent Ethanol Exposure Impacts Offspring Transcriptomes with Sex and Tissue Specific Effects"

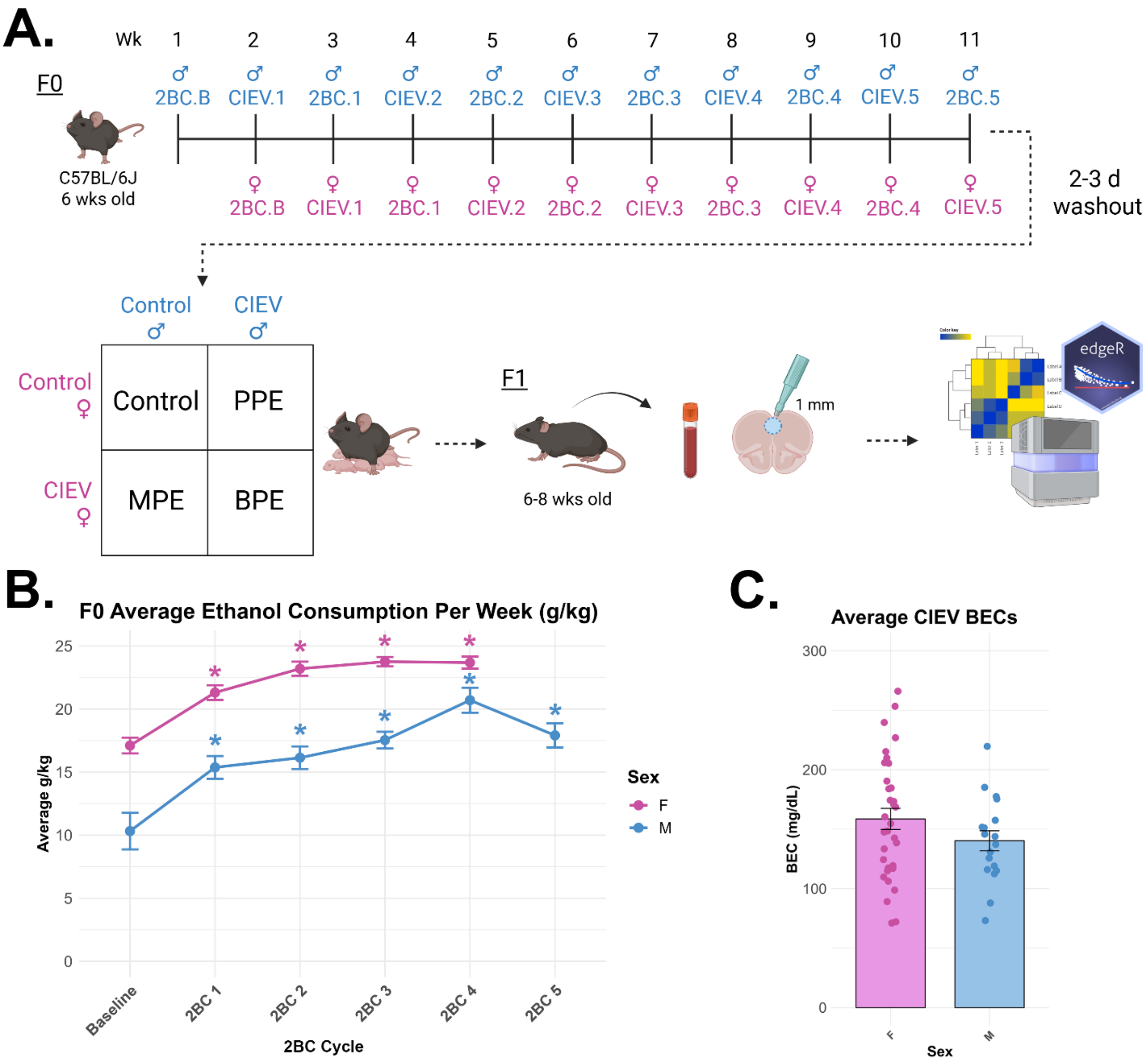


**Supplementary Figure S1. Whole blood and mPFC study overview and CIEV-2BC validation.** (A). Graphical study methods. Male and female founder mice (F0 generation) were interleaved in five cycles of CIEV-2BC. Following a 2–3-day washout period after the last ethanol exposure, mice were randomly mated to produce PPE, MPE, and BPE offspring (F1 generation). Adult, ethanol-naïve F1 mice were sacrificed in adulthood (6-8 weeks of age) and whole blood as well as 1-mm mPFC tissue punches were taken for RNA-Sequencing. (B.) Ethanol consumption (in g/kg) in males and females by 2BC cycle. Data are presented as mean ± SEM. A linear mixed-effects model revealed significant main effects of Sex, $F(1, 52.35) = 71.82$, $p < .001$, and Week, $F(5, 214.29) = 44.48$, $p < .001$, as well as a significant Sex × Week interaction, $F(4, 214.29) = 3.46$, $p = .009$. Post-hoc pairwise comparisons (Tukey-adjusted) revealed significantly increased ethanol consumption from baseline in both males and females for all cycles of 2BC ($p < 0.001$ for all weekly comparisons to baseline within sexes). (C.) BECs achieved during CIEV exposure, merged across cycles. Data are presented as mean ±SEM. *Panel A was created using Biorender.com.*

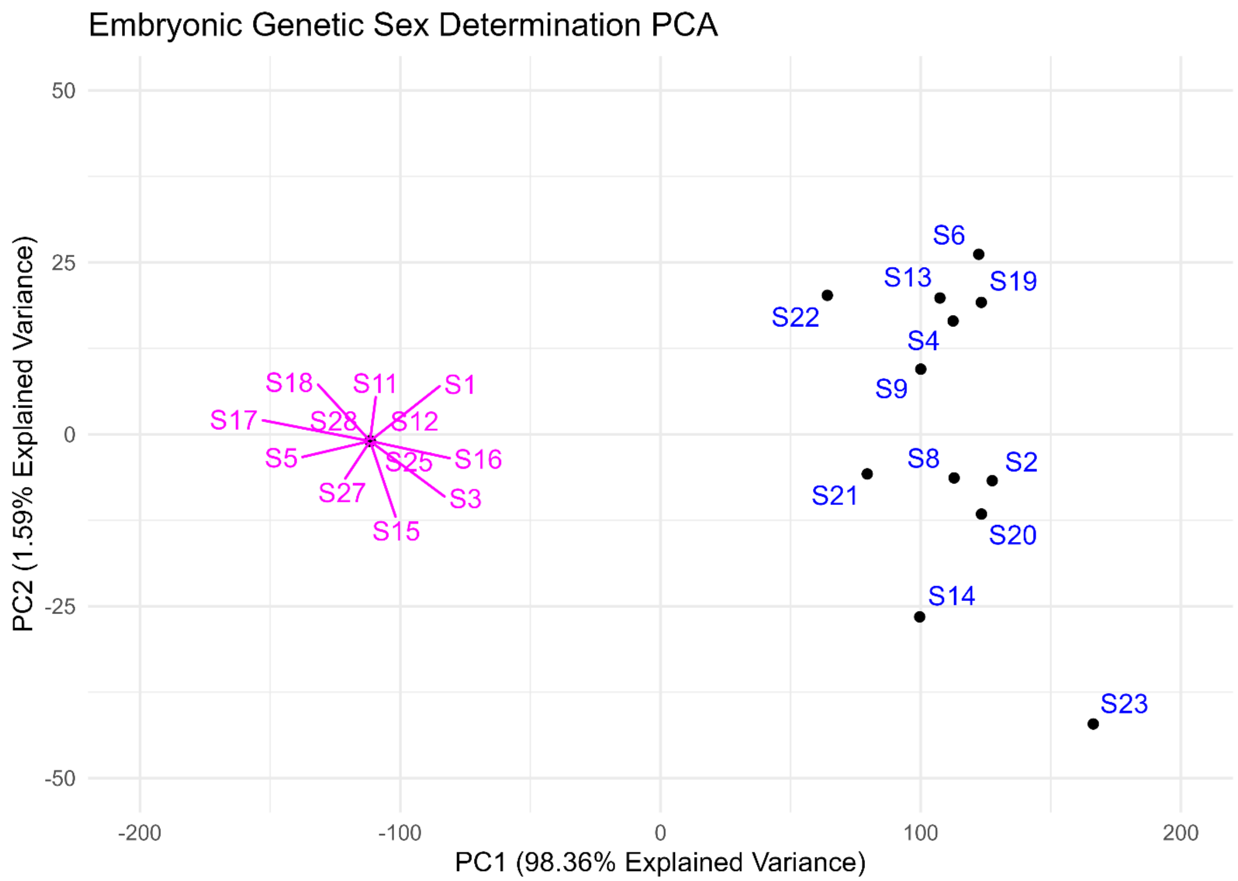


**Supplementary Figure S2. Principal component analysis (PCA) to determine genetic sex of PPE embryos.** PCA was performed on the counts per million (CPM) values of the Y chromosome genes *Ddx3y*, *Kdm5d*, and *Eif2s3y*. The resulting biplot of principal components one and two (PCA2 vs PCA1) is shown. PPE embryos designated female formed a tight cluster with near-zero CPM values, while PPE embryos designated male formed a cluster with appreciable expression levels of *Ddx3y*, *Kdm5d*, and *Eif2s3y.*
